# Chromosome specific telomere lengths and the minimal functional telomere revealed by nanopore sequencing

**DOI:** 10.1101/2021.06.07.447263

**Authors:** Samantha L. Sholes, Kayarash Karimian, Ariel Gershman, Thomas J. Kelly, Winston Timp, Carol W. Greider

## Abstract

We developed a method to tag telomeres and measure telomere length by nanopore sequencing in the yeast *S. cerevisiae.* Nanopore allows long read sequencing through the telomere, subtelomere and into unique chromosomal sequence, enabling assignment of telomere length to a specific chromosome end. We observed chromosome end specific telomere lengths that were stable over 120 cell divisions. These stable chromosome specific telomere lengths may be explained by stochastic clonal variation or may represent a new biological mechanism that maintains equilibrium unique to each chromosomes end. We examined the role of *RIF1* and *TEL1* in telomere length regulation and found that *TEL1* is epistatic to *RIF1* at most telomeres, consistent with the literature. However, at telomeres that lack subtelomeric Y’ sequences, *tel1Δ rif1Δ* double mutants had a very small, but significant, increase in telomere length compared to the *tel1Δ* single mutant, suggesting an influence of Y’ elements on telomere length regulation. We sequenced telomeres in a telomerase-null mutant (*est2Δ*) and found the minimal telomere length to be around 75bp. In these *est2Δ* mutants there were many apparent telomere recombination events at individual telomeres before the generation of survivors, and these events were significantly reduced in *est2Δ rad52Δ* double mutants. The rate of telomere shortening in the absence of telomerase was similar across all chromosome ends at about 5 bp per generation. This new method gives quantitative, high resolution telomere length measurement at each individual chromosome end, suggests possible new biological mechanisms regulating telomere length, and provides capability to test new hypotheses.

## INTRODUCTION

Maintenance of the telomere length equilibrium is critical for cell survival. Telomeres shorten with each cell division while a few are stochastically elongated by the enzyme telomerase (Greider and Blackburn 1985; Teixeira et al. 2004). This balance of shortening and lengthening is exquisitely regulated to generate a normal distribution of lengths that is established around a mean or “set point”. Different species have very different telomere length set points; for example *Tetrahymena* has 70 bp, yeast have 300 bp, humans have 10kb, and some mouse strains have up to 80kb mean telomere lengths (Greider 1996). How this distribution is established and maintained is not fully understood. When telomerase levels are low or absent, the distribution shifts toward shorter lengths and the shortest telomeres signal a DNA damage response. These cells undergo a permanent cell cycle arrest, termed senescence or apoptosis, depending on the cell type (Enomoto et al. 2002; d’Adda di Fagagna et al. 2003; IJpma and Greider 2003).

Loss of telomere length maintenance in humans leads to disease. Individuals with inherited mutations in telomerase, or other telomere regulatory genes, have Short Telomere Syndrome, which includes bone marrow failure, immunodeficiency, pulmonary fibrosis and other diseases (Armanios 2013). In contrast, activation of telomerase (Counter et al. 1992; Greider 1998) or inheritance of long telomeres, allows cell immortalization which can predispose to cancer (Stanley and Armanios 2015; McNally et al. 2019). Thus, deviation from the mean telomere length in either direction affects cellular lifespan and plays a critical role in disease.

The yeast *S. cerevisiae* serves as an excellent model for probing mechanisms of telomere length regulation. Many genes and regulatory mechanisms that affect human telomere length were first discovered in yeast (Wellinger and Zakian 2012). Yeast telomere sequences are more variable than the consistent TTAGGG_n_ repeats found in all vertebrates (Moyzis et al. 1988). They are composed of a mixture of GT, GGT or GGGT sequence motifs, usually abbreviated, G_1-3_T. The subtelomeric region in yeast contains blocks of repeated DNA.

Deletion of any component of telomerase in yeast leads to progressive telomere shortening, termed ever-shorter telomere (EST) phenotype (Lundblad and Szostak 1989; Lendvay et al. 1996) Previous work suggests that telomeres shorten by about 3-5 bp per cell division in the absence of telomerase (Marcand et al. 1999; Wellinger and Zakian 2012; Xu et al. 2013). There is initially no effect of telomere shortening on cell growth and survival, but when telomeres become very short after passaging in culture for several days, most cells arrest at G2/M (Enomoto et al. 2002; IJpma and Greider 2003). A few cells can rescue their short telomeres through recombination and will generate “survivors” that grow in the absence of telomerase (Lundblad and Blackburn 1993). Survivors are generated through a break induced replication (BIR) pathway (Bosco and Haber 1998; McEachern and Haber 2006) that can initiate within the telomere repeats (Type I survivors) or in the subtelomere elements (Type II survivors) (Teng and Zakian 1999). Deletion of the recombination gene *RAD52* significantly reduces production of survivors (Lundblad and Blackburn 1993) although very rare survivors have been found in *est2Δ rad52Δ* double mutants (Lebel et al. 2009).

The normal distribution of the telomere length equilibrium is due primarily to sequence loss through replication and sequence addition by telomerase. It has been proposed that each of the 32 chromosome ends has a similar distribution of telomere lengths (Shampay and Blackburn 1988). Shampay and Blackburn showed that each end can slightly shift its mean length due to clonal variability. If a cell that happens to have a short telomere at chromosome 8L, for example, seeds a new colony on a plate, the descendants of that cell will initially have a new shorter telomere length distribution until enough divisions occur in the population to homogenize all ends towards a population mean (Shampay and Blackburn 1988).

Telomerase adds between 3bp to 179 bp onto chromosome ends in one cell cycle, however only a few telomeres are elongated at each cycle (Teixeira et al. 2004). Sequence analysis suggests that short telomeres are preferentially elongated compared to long telomeres (Teixeira et al. 2004). There is also evidence that replication fork collapse in telomere repeats, followed by telomerase addition, may contribute to the appearance that short telomeres are preferentially elongated (Paschini et al. 2020). How preferential elongation of short telomeres or replication fork collapse allows the robust maintenance of the telomere length equilibrium that is observed over many cell divisions is not yet clear.

The standard technique for visualizing telomere length in yeast is by Southern blot. Genomic DNA from yeast cells is cut with a restriction enzyme near the telomere and hybridized with a probe to the telomere repeats or to the subtelomere, generating smeared bands visible on the autoradiogram (Shampay and Blackburn 1988). The smears represent the heterogenous distribution of telomere lengths across both the different chromosomes and between cells. Since restriction sites on different chromosomes are at different locations, the absolute size of each smear differs. Frequently, the restriction enzyme XhoI is used to cut genomic DNA because many telomeres have a conserved XhoI restriction site in the Y’ subtelomere element. Multiple telomeres are then visualized in a heterogeneous band centered around 1.2Kb. Measuring median length of this 1.2kb sized band can allow large changes in bulk telomere length to be measured in various mutant backgrounds, but it is not possible to deconvolute the length of individual telomeres from each other.

We developed a nanopore sequencing method to measure telomere length to provide a more quantitative analysis of telomere length variation. Nanopore sequencing allowed us to capture a full telomere sequence as well as the subtelomere and unique chromosomal sequence in one long read. This provides quantitative measurement at the single molecule of the telomere length distribution. Using this method, we found each unique chromosome end has its own specific length distribution and, surprisingly, this distribution is maintained for hundreds of generations. We also found end specific effects of the length regulators *TEL1* and *RIF1.* In the absence of telomerase, all chromosomes shortened at a rate of 5pb per generation, and there were frequent telomere recombination events before bulk survivors appeared in the population. Combining data from all of these experiments, we estimate that the minimal length for telomere function in yeast is 75 bp. Identifying these nuanced changes with the increased resolution and precision of nanopore sequencing will allow for additional mechanistic insights into telomere length regulation.

## RESULTS

### Telomere tagging and bioinformatic analysis

To accurately measure telomere length and retain the telomeric 3’ overhangs, we modified a method (Teixeira et al. 2004) to tag the molecular end of the chromosome. We prepared high molecular weight DNA(Denis 2018) and added poly(A) to 3’ ends with terminal transferase. We next annealed an oligo dT primer that also contained a unique adapter sequence, termed TeloTag. Addition of Sofobulus DNA polymerase IV, which lacks strand displacement and exonuclease activity, was used to fill in the opposite strand, and T4 DNA ligase to seal the nicks. Nanopore library adaptors were added to allow sequencing (Fig. 1A).

**Figure 1.**
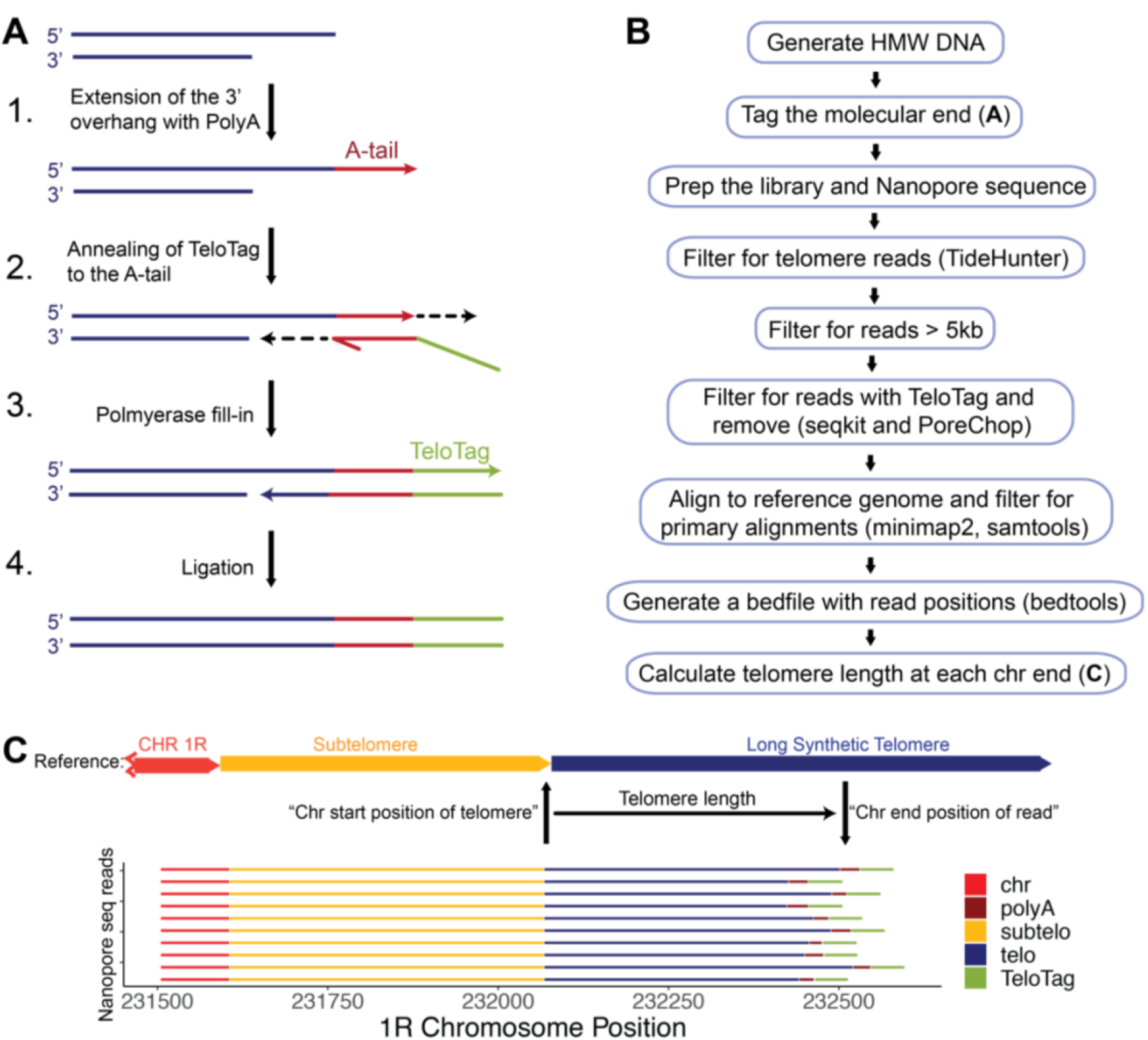
Telomere length measurement by nanopore sequencing. A) Schematic of method to tag the molecular end of the chromosome. 1. Terminal transferase adds poly A sequence to the 3’ overhang of the telomere. 2. A complementary oligo-dT sequence on the TeloTag oligonucleotide end containing a unique TeloTag sequence is annealed to the poly(A) sequence. 3. Polymerase fill-in generates double stranded blunt ends. 4. Ligase is used to seal the nick. Nanopore adaptors are then added using the standard library prep. B) Bioinformatic pipeline to generate telomere length information (see methods). C) Telomere length is calculated using the chromosomal position of each telomere sequence start and calculating the distance to the TeloTag sequence at the chromosome end. A random sample of 10 sequence reads aligned at CHR 1R are presented as an example.

Tagged genomic DNA was run on nanopore MinIon or Flongle flow cells to generate Whole Genome Sequencing (WGS) data. For optimal use of a nanopore MinIon flow cell, samples were multiplexed, and only reads with barcodes on both ends were retained. Reads with telomeric sequence were extracted using TideHunter and retained only if they contained the TeloTag and were longer than 5 kb to ensure that they could be mapped through the telomere and subtelomere into the unique chromosomal regions (Gao et al. 2019). Adapter sequences were then bioinformatically removed using Porechop (Wick 2018). These reads were aligned using minimap2 to a *S. cerevisiae* reference genome, sacCer3 (Foury et al. 1998)genome onto which we added 2kb of synthetic telomere sequence to ensure that the longest telomeres were captured (see Methods). Finally, the unique telomere start sequence, the position where the subtelomere ends and telomere repeats began, was identified for each chromosome end. The telomere length was calculated by measuring the distance from the chromosomal position where the telomere sequence started to the position where the aligned telomere read ended (Fig. 1C).

### Telomere length by nanopore sequencing recapitulates length measurement by Southern blot

To test the accuracy of our nanopore method for telomere length measurement, we compared telomere fragment length analysis by nanopore and Southern blot. We first looked at strains with expected clear differences in telomere length profiles, wildtype and *rif1Δ.* Deletion of the *RIF1* gene results in very long telomeres on Southern blots (Hardy et al. 1992), allowing us to probe a greater distribution of telomere length profiles and test whether these length differences are captured with nanopore. We measured telomere fragment length after XhoI digest using a probe against the common Y’ subtelomere element; this will measure telomere lengths in bulk, probing 17 of 32 subtelomeres (Fig. 2A, Supplemental Fig. S1C). We ran all samples in duplicate on a Southern blot and included an additional probe to the unique sequence at CEN4 that runs at 1.4kb as an internal molecular marker for densitometry plots (Fig. 2A, Supplemental Fig. S1C). We used densitometry scanning to quantify the signal on the Southern blot (see Methods).

**Figure 2.**
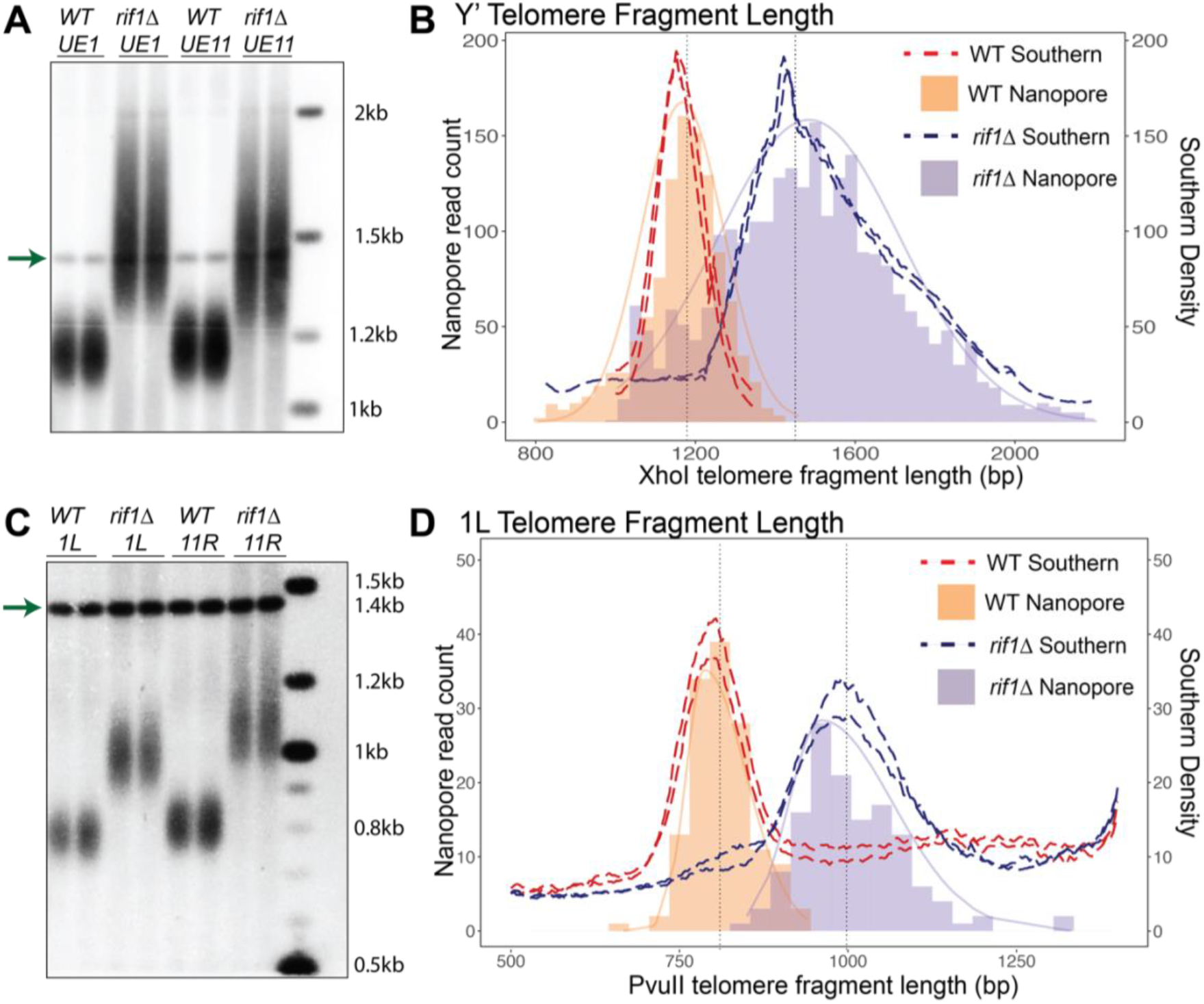
Telomere fragment length measured by Nanopore sequencing accurately reflects length measurements by Southern blot. A) Southern blot probed with Y’ subtelomeric probe and with CEN4 probe (arrow) as a marker at 1.4kb to calibrate for densitometry. Two replicates of wildtype (WT) cells and two replicates of *rif1Δ* cells are shown for each strain, UE1 and UE11. B) Comparison of densitometry (dotted lines) of Southern blot shown in A) and Nanopore sequencing (histogram fill bars) for UE1 strain. The light orange bars represent nanopore read counts for WT cells, the light purple bars represent nanopore read counts for *rif1Δ* cells. C) Southern blot probed with the Unique single telomere 1L probe and CEN4 probe (arrow). D) Comparison of densitometry (dotted lines) of Southern blot in C) and Nanopore sequencing (bars) of UE1 1L telomeres. The light orange bars represent nanopore read counts for WT cells, the light purple bars represent nanopore read counts for *rif1 Δ* cells.

On the same samples, we performed our nanopore sequencing method as described (Fig. 1; Methods). Nanopore sequencing data statistics are summarized in Supplemental Table S1. To directly compare to Southern blot telomere length, we included the internal chromosomal sequence up to the XhoI site for Y’ on the nanopore telomere read file. The histogram of the nanopore telomere sequence read lengths aligned remarkably well with the Southern blot densitometry for both wildtype telomeres and the much longer *rif1Δ* telomeres in two different strains (Fig. 2B, Supplemental Fig. S2A). We used WALTER, a tool for Southern blot densitometry, that generates summary length statistics (Lycka et al. 2021), to compare Southern blot and nanopore telomere length profiles, and again saw comparable profiles (Supplemental Fig. S3A). Since telomere length profile is predicted to be a normal distribution, we generated Q-Q plots to examine the distribution of nanopore and Southern density data against a normal distribution (Supplemental Fig. S4A&B) and against each other (Supplemental Fig. S4C). The linearity of the Q-Q plots of both the Southern and nanopore data confirm that these profiles are a normal distribution (See Supplemental Fig S4 for details)

Because the Y’ probe measures 17 telomere length distributions simultaneously, it is challenging to detect potential small effects and chromosome specific telomere lengths with this probe on a Southern blot. To assess the length distribution of single chromosome ends, we generated strains where the subtelomere was replaced by a unique sequence. We used two different strains (Shubin et al. 2021) with unique sequence at either chromosome end 1L Unique End 1 (strain UE1), or at chromosome end 11R: Unique End 11 (strain UE11) (Supplemental Fig. 1A/ B). We examined the telomere lengths of these single 1L and 11R chromosome ends in both wildtype and *rif1Δ* backgrounds, using a probe to the unique ends (Fig. 2C, Supplemental Fig. S1D). Comparing these single telomere Southerns to the nanopore sequencing data we again saw very high agreement for both wildtype and *rif1Δ* telomere lengths (Fig. 2D, Supplemental Fig. 2B). Southern blot WALTER densitometry profiles (Lycka et al. 2021) versus the nanopore data also showed similar profiles between Southern density distribution and the nanopore length distribution (Supplemental Fig. S3B), suggesting that nanopore sequencing can accurately measure telomere fragment length in a bulk population or single telomere population across different telomere length profiles.

To determine the minimum number of reads required for telomere length measurement using nanopore sequencing, we compared the MinIon flowcells to the lower cost, but lower yield, Flongle flowcells. Multiplexing two samples per Flongle and MinIon flowcell yielded many fewer reads from the Flongle (~150-250 per barcode) vs the MinIon (~2000 reads per barcode), but we found no statistically significant differences between the mean and distributions of telomere length across technical replicates and flow cell types (Supplemental Fig. S5). We conclude that the telomere length assay can either be highly multiplexed or can generate data on a lower yield flowcell to easily generate a bulk telomere measurement.

### Each chromosome end shows reproducible chromosome specific telomere length distributions

Probing telomere lengths on unique chromosome ends is challenging with Southern blots, but nanopore sequence reads are long enough to span the subtelomeric elements and extend into unique chromosomal sequence. To examine individual chromosome ends, we sequenced three independent wildtype clones and three *rif1Δ* clones (Supplemental Fig. S6B). For the wildtype, we used two independent clones of UE1 and one clone of UE11. Each clone was isolated by picking an independent single colony from a freshly streaked plate of cells (Supplemental Fig. 6B). We analyzed both the bulk telomere length of all telomeres and individual chromosome ends (Fig. 3A/B).

**Figure 3.**
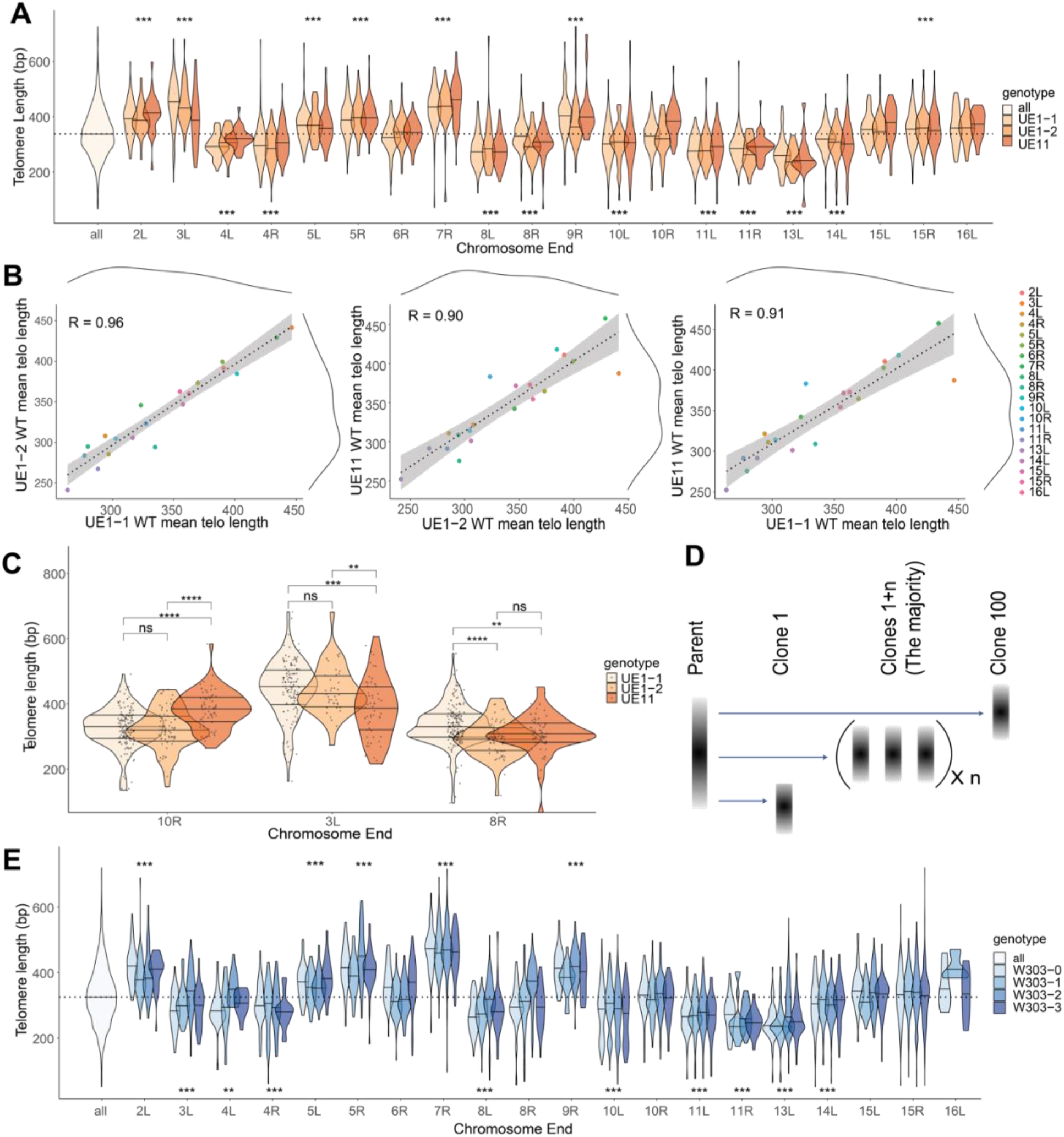
Nanopore sequencing mapping of the telomere length distribution of each individual chromosome end. A) Bulk telomere length (all) and telomere length for each chromosome end were mapped by aligning to internal unique sequence for WT cells. Three independent clones were sequenced two from UE1 (UE1-1 and UE1-2) and one from UE11 (see supplemental figure S6). Telomere lengths are reported as violin plots with a line at the mean. A dotted line shows the mean of the total population at 343.3 for WT cells. Analysis of the means (ANOM) multiple contrast test of each telomere against the grand mean of all telomere length profiles was performed (see Supplemental Figure S7A) and p values adjusted by the Bonferroni method are reported as *<0.05, **<0.01, ***<0.001, ****<0.0001, or not significant (n.s.). B) Pairwise correlations of mean telomere length across each of the three replicates for WT. A linear model was fit to the data (dotted line) to calculate the correlation constant, reported as the R^2^ value. Marginal density of each dataset is plotted as solid black lines across the top and right of the plots. C) Clonal variation was observed at 10R, 3L, and 8R in WT cells. Individual telomere length reads are represented as single points within the violin plots. The significance of the difference between two sample was tested using a multiple t-test (methods) and p values adjusted by the Bonferroni method are reported as above. D) Model of clonal variation where the telomere length of the majority of clones remains at the parental average, but occasional length outliers may seed new distributions represented by Clone 1 and Clone 100. E) Telomeres for each chromosome end were mapped by aligning to internal unique sequence. Four independent clones were sequenced from the WT W303-0 parent strains of UE1 and UE11 (See Supplemental Figure S8A). Analysis of the means (ANOM) multiple contrast test of each telomere against the grand mean of all telomere length profiles was performed (see Supplemental Figure S7B) and p values adjusted as stated above.

We were able to individually map 23 of 32 total chromosome ends with unique subtelomere and upstream genic sequences. However we were not able to map 9 chromosome ends uniquely (2R, 3R, 6L, 9L, 12L, 12R, 13R, 14R, 16R) due to variation in subtelomeric structure from the reference genome. Subtelomeric elements are known to recombine and generate end diversity (Louis and Haber 1990; Louis and Haber 1992; Maxwell et al. 2004). The reference genome we used to map the subtelomere sequences (s288C) likely has a different arrangement of subtelomeric elements than the strain we were sequencing (w303). We thus excluded these difficult to map telomeres in our analysis.

We were surprised to find that some chromosome ends showed telomere length distributions that were consistent across three independent clones, but distinctly longer or shorter from the population mean of all telomeres (Fig. 3A). To quantify the differences, we applied analysis of the means (ANOM) to compare the mean length of individual chromosome ends to the population mean length of all the telomeres, and found that eight chromosome ends (4L, 4R, 8R, 10L, 11L, 11R, 13L, and 14L) had shorter telomeres than the population mean, while six chromosome ends (2L, 3L, 5L, 5R, 7R, and 9R) had longer telomeres than the population mean (Supplemental Fig. S7A). A pairwise comparison of the three datasets suggests that telomere length of individual chromosome ends was consistent across replicates with correlation coefficients (R) of 0.96, 0.90, and 0.91(Fig. 3B.) We note that for chromosomes where both ends are analyzed, there was a trend toward concordance with both ends being long like 5L/5R, or both ends being short like 4L/4R 8L/8R and 11L/11R. While we demonstrated earlier that wildtype telomere length distribution is normal by Q-Q plot, *rif1 Δ* telomere length did not appear to be maintained in a Gaussian distribution (data not shown). In the *rif1Δ* mutant, we saw a lack of correlation between individual chromosome ends between replicates (Supplemental Figure S9). The disruption of these chromosome end-specific telomere lengths in *rif1Δ* cells suggests that these differences are not detectable due to greater heterogeneity in the length distribution or to loss of telomere length control in the mutant, or both.

The apparent chromosome end-specific telomere lengths in wildtype cells could be due to stochastic clonal variation as previously proposed (Shampay and Blackburn 1988). That is, when a cell with a short telomere on chromosome 4L forms a new colony, it will seed a new telomere distribution that will center around a shorter midpoint (Fig. 3D). We found statistically significant differences at 3L, 8R, and 10R across the three clones, at 3L, UE11 was significantly shorter than UE1-1 and UE1-2, at 8R UE1-1 was significantly longer than UE-1 and UE11, and at 10R UE1-1 and UE1-2 were significantly different than UE11. We were surprised that clonal variation was not more frequent given the apparent frequent occurrence of clonal variation in the previous study (Shampay and Blackburn 1988), although this may be due to different number of cell divisions in propagating different clones.

Shampay et al. introduced the concept of the telomere length ‘set point’, in which all telomeres will gradually move back toward a population mean length distribution. This suggests that given enough cell divisions in the presence of telomerase, we would expect all chromosome ends to approach a mean value. In fact, Shampay et al. showed that continued growth in liquid culture led to a broadening of the length distribution, which they interpreted as homogenization of all end lengths (Shampay and Blackburn 1988). This combined with other observations (Marcand et al. 1997) has led to establishment of a “protein counting” model, where telomerase is proposed to preferentially elongate short telomeres while ignoring long ones. These models predict homogenization of telomere length after many opportunities for telomere shortening and lengthening. To test these models and see whether the lengths would homogenize with more divisions, we grew cells for 120 generations and sequenced three independent clones (Supplemental Figure S8A). We sequenced W303-0, the parent strain (derived from CVy61, see Methods) that gave rise to both UE1 and UE11 (Supplemental Fig. S6A). We passaged the W303-0 cells in liquid culture for approximately 120 cell doublings, then plated and selected three independents colonies for sequencing (W303-1, W303-2, and W303-3) (Supplemental Fig. S8A). We expected, based on previous models, that the telomere length distributions at individual chromosome ends would revert towards a grand population mean length with more cell divisions. Surprisingly, we saw that some chromosome specific telomere lengths were maintained across all W303 clones, for example: 2L, 5L, 5R, 7R, 8L, 9R, 10L, 11L, 11R, and 13L (Fig. 3E, Supplemental Fig. S7B). Although there were some examples of large deviations from the initial length, for example at 3L, 6R, 15L and 16L, the major trend was reproducibility of the parents’ chromosome specific lengths (Supplemental Fig. S7A and B). To quantify the variation between different clones, we examined pairwise correlations between the W303-0 parent and the subclones, and found correlation coefficient values (R) of 0.92, 0.81, 0.98, 0.87, 0.90, and 0.84 (Supplemental Fig. S8B). Our data indicate that after 120 doublings, the chromosome end specific telomere lengths are largely maintained. This implies a very slow rate of reversion to a common mean length distribution, if it occurs at all, and implies new models for telomere length regulation are needed to explain this data.

### Subtelomeric Y’ elements have a small effect on telomere length regulation by *TEL1* and *RIF1*

The possibility of chromosome end-specific telomere lengths raised the idea that subtelomeric sequence might influence telomere length. A similar suggestion was made by Craven and Petes who proposed that the subtelomeric X and Y’ elements influenced the effects of *TEL1* and *RIF1* on telomere length (Craven and Petes 1999). Specifically, they suggested that *TEL1* is epistatic to *RIF1* only at telomeres with Y’ elements, but not at telomeres with only X elements. That is, telomeres are long in *rif1Δ* mutants, short in *tel1Δ* mutants, and also short in the *tel1Δ rif1Δ* double mutant at Y’ telomeres.

We reexamined this question using both Southern blots and nanopore sequencing. We generated *tel1Δ*, *rif1Δ,* and *tel1Δ rif1Δ* mutants in the in UE1 background. As expected, in the Southern blot probed with Y’ sequence *rif1Δ* mutants had long telomeres and *tel1Δ* mutants had short telomeres. The double mutant, *tel1Δ rif1Δ,* had short telomeres very similar to the single *tel1Δ* mutant (Fig. 4A), indicating that *TEL1* is epistatic to *RIF1* at telomeres with Y’ elements. Nanopore sequencing showed the same result at chromosomes with Y’ elements; *tel1Δ rif11Δ* double mutants had short telomeres indistinguishable from *tel1Δ* telomere lengths (with the exception of 14L), supporting the previous finding (Fig. 4B). However, for most chromosomes ends that had only X elements, telomere length in *tel11Δ rif11Δ* was slightly longer (~65bp on average) and statistically different from *tel1Δ* (Fig. 4D). This trend also was seen at the unique chromosome 1L telomere where the subtelomeric sequence was removed entirely. Both nanopore sequencing (Fig. 4D, 1L) and Southern blot (Fig. 4C) showed that telomeres in the *tel1Δ rif1Δ* double mutant were longer than the *tel1Δ* alone. While the difference in length was not as large as seen by Craven and Petes, it is reproducible and suggests interesting unexplored biology.

**Figure 4.**
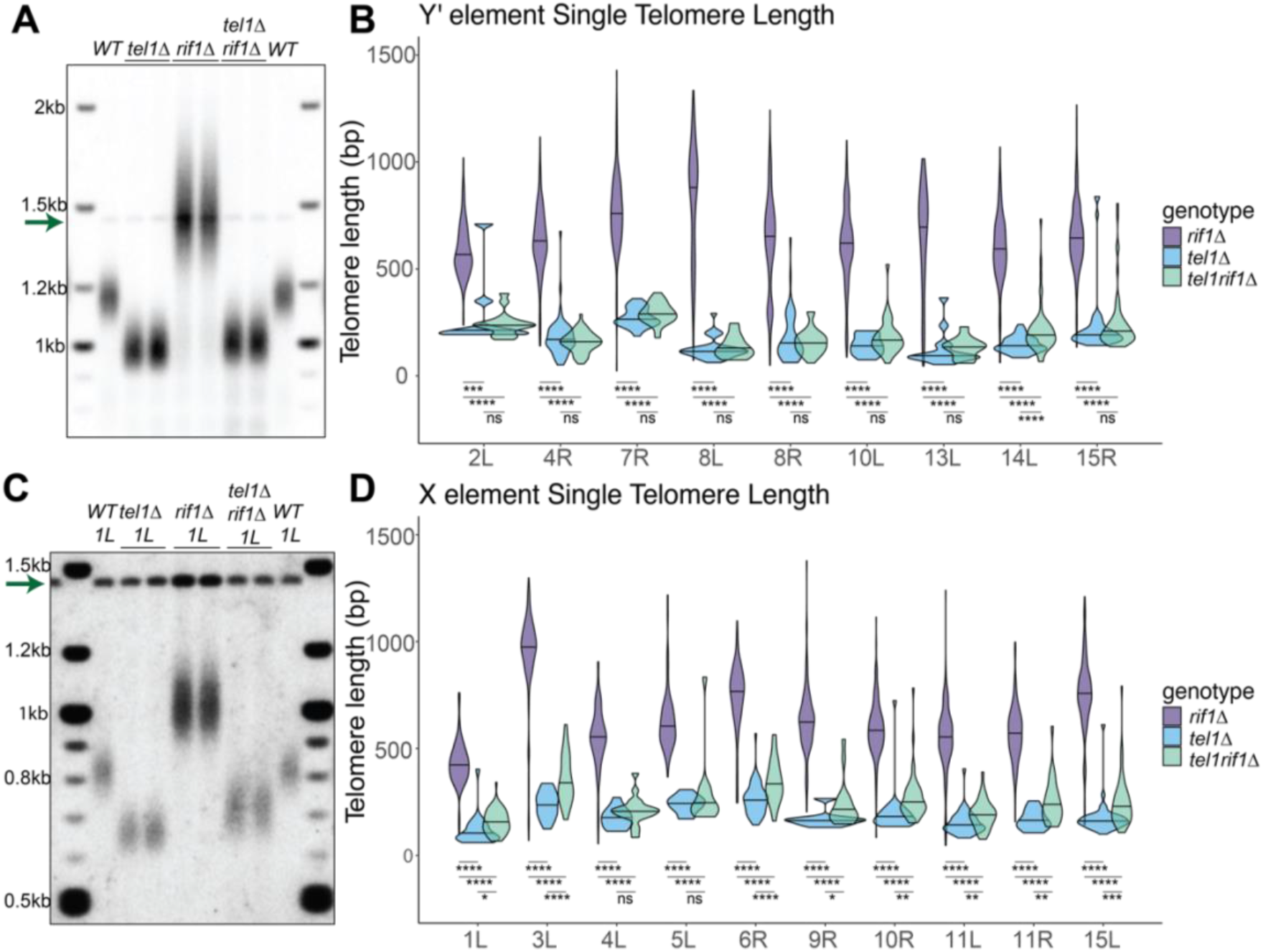
Subtelomeric sequences have a small effect on telomere length regulation by *TEL1* and *RIF1*. A) Southern blot of WT, *tel1Δ, rif1Δ,* and *tel1Δ rif1Δ* cells with bulk telomere length hybridized with the Y’ and CEN4 probes (arrow). B) Individual telomere lengths determined by nanopore for telomere ends containing Y’ subtelomeric elements reported for *tel1Δ, rif1Δ,* and *tel1Δ rif1Δ* cells. C) Southern blot of WT, *tel1Δ, rif1Δ, and tel1Δ rif1Δ* samples hybridized with Unique probe to 1L and CEN4 probe (arrow) to examine unique telomere 1L with no subtelomeric elements. D) Individual telomere lengths determined by nanopore for telomere ends containing only X subtelomeric elements or no telomeric elements (1L) reported for *tel1Δ, rif1Δ,* and *tel1Δ rif1Δ* cells. A multiple t-test was used to assess the significance of telomere length differences between genotypes, and p values are reported as *<0.05, **<0.01, ***<0.001, and ****<0.0001, or not significant (n.s.).

### Telomere shortening in the absence of telomerase reveals the minimal functional telomere

To examine the dynamics of telomere loss in cells without telomerase, we examined telomerase null cells. We sporulated an *EST2/est2Δ RAD52/rad52Δ* heterozygous diploid and selected individual haploid colonies of *est2Δ, est2Δ rad52Δ,* and wildtype (*EST2*). We grew these genotypes from a single colony and passaged each three times in liquid culture (Supplemental Fig. S10). To determine cell division numbers, we estimated growth from a single cell to colony formation to be about 20 cell doublings. Both *est2Δ* and *est2Δ rad52Δ* cells grew slowly after 55 population doublings, and *est2Δ rad52Δ* cells exhibited significantly slower growth than *est2Δ.* Telomere shortening in the absence of telomerase leads to cell cycle arrest (Enomoto et al. 2002; IJpma and Greider 2003)), and also to telomeric elongation by Break Induced Replication (BIR) (Bosco and Haber 1998) that ultimately allows the outgrowth of “survivors” (Lundblad and Blackburn 1993). This implies there must be a competition between arrest and BIR, and presumably all chromosomes must be elongated for phenotypic survivors to emerge.

Telomere lengths in survivors appear on Southern blots as substantial increases likely because a significant number of ends must elongate to overcome senescence and to visualize these events on a Southern. With nanopore we saw moderate increases at single telomeres that are not visible in the bulk population and would not be possible to detect by Southern (Fig. 5A/B). For example 1L, 2R, 3L, 4L, 8R, 9R, 10R, 11R, 13R, and 15R all show long telomere outliers (Fig. 5C). In some cases these outliers are seen in the first passage for example 1L and 10R, while in other cases they appear in the later passages as for 11R. In the absence of telomerase, we expect that this generation and loss of these longer telomeres will be stochastic and any given cell with an elongated chromosome end; for example 1L might senesce and not generate a survivor due to shortening of some other chromosome end.

**Figure 5.**
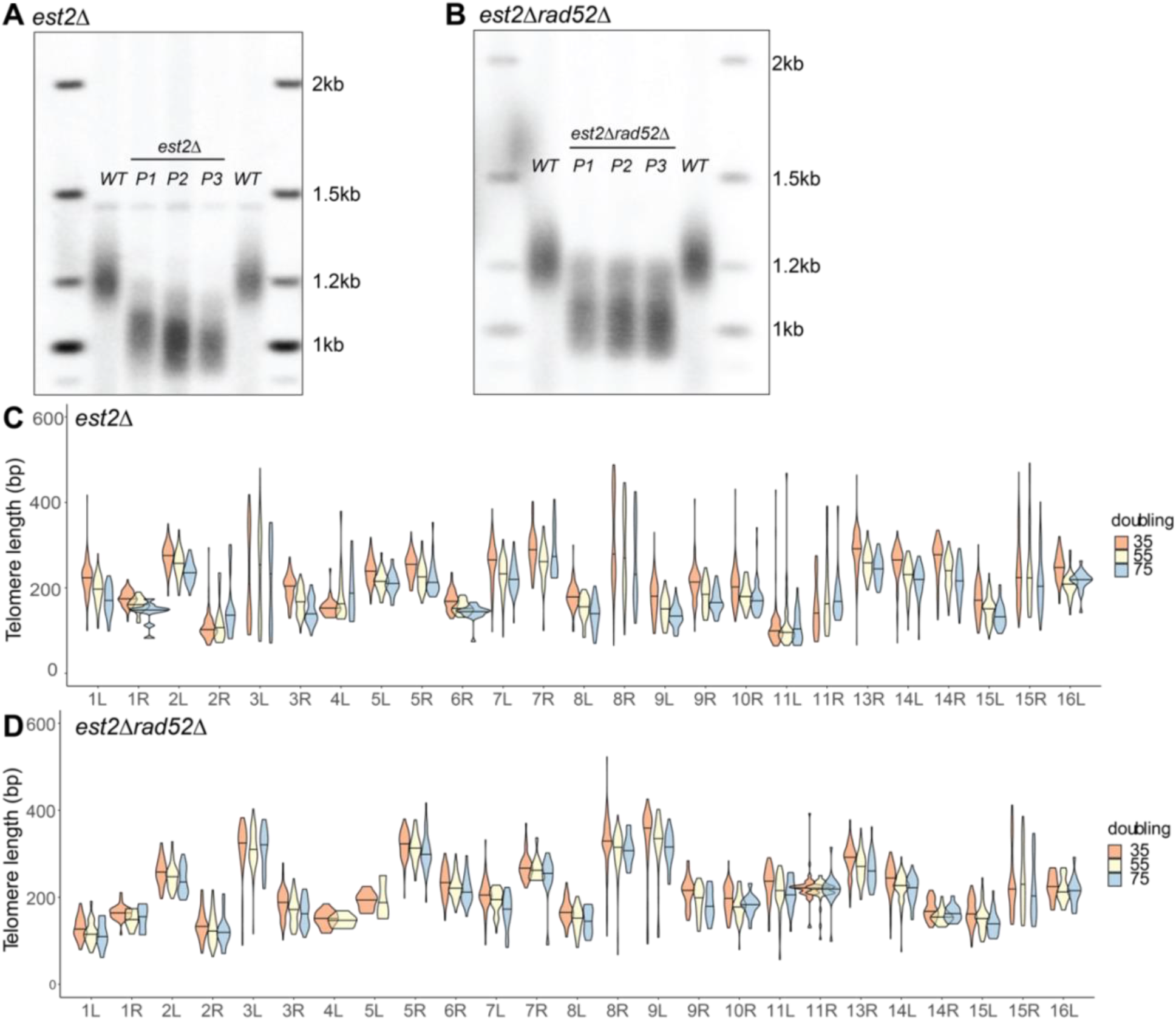
Telomere shortening in a telomerase-null, *est2Δ*, background at individual telomeres over time. Freshly generated *est2Δ* and *est2Δ rad52Δ* haploids were grown up (P1, 35 doublings) and passaged twice (P2, 55 doublings and P3, 75 doublings) to allow for telomere shortening (see Methods). Telomere length was measured by Southern blot hybridized with the Y’ probe and by nanopore sequencing on the same samples. A) Southern blot of WT, P1, P2, and P3 *est2Δ* hybridized with the Y’ probe. B) Southern blot of WT, P1, P2, and P3 *est2Δ rad52Δ* hybridized with the Y’ probe. C) Individual telomere length of *est2Δ* cells determined by nanopore sequencing of P1, P2, and P3. D) Individual telomere length of *est2Δ rad52Δ* cells determined by nanopore sequencing of P1, P2, and P3.

To further examine the role of BIR in the generation of outliers, we examined the *est2Δ rad52Δ* cells, which are recombination deficient. If the long telomere outliers in *est2Δ* cells are due to BIR, we would expect to see fewer of these length increases in *est2Δ* rad52*Δ* cells. Indeed, we saw only two examples of very long outlier telomeres in the *est2Δ rad52Δ* cells compared to twelve *est2Δ* cells. (Fig. 5D). We did see outliers at chromosome 8R and 11R in *est2Δ rad52Δ.* These length increases may be due to rare events that occur even in *est2Δ rad52Δ* cells that allow telomere elongation (Lebel et al. 2009; Claussin and Chang 2016).

To determine the rate of telomere shortening in the absence of telomerase before cell division stops, we compared the rate shortening at individual chromosome ends in *est2Δ* and *est2Δ rad52Δ* cells over three outgrowths or passages (P1, 2, 3) (Supplemental Fig. S10). We exclude data with apparent recombination events, as described above, as it would skew the analysis. To establish the initial rate of shortening, we used the telomere length for each chromosome from the wildtype parent strain and termed this P0 because the initial *est2Δ* haploid colony already had undergone 20 cell division in the absence of telomerase (Supplemental Fig. S10). We note that that these P0 values are only estimates because they are not from the same *est2Δ* passaged cells, though our earlier data suggests that telomere lengths remain consistent at individual chromosome ends. We then also plotted the rates from P1 to P3 and found different slopes; the first from P0 to P1, where few cells may experience critically short telomeres, was 5.33 bp/cell division in *est2Δ* and 5.13 bp/cell in *est2Δ rad52Δ* cells (Fig. 6A/B). The second rate, from P1 to P3, showed a slope of 0.92 bp/cell division in *est2Δ* cells and a slope of 0.39 bp/cell in *est2Δ rad52Δ* cells (Fig. 6A/B). This telomere shortening rate is similar to other estimates of 3-4 bp/generation (Marcand et al. 1999; Abdallah et al. 2009; Wellinger and Zakian 2012; Xu et al. 2013). The second slope could be less steep because cells with very short telomeres don’t divide and thus are lost from the calculated rate. The slopes of the P1 to P3 *est2Δ* cells were higher than those of *est2Δ rad52Δ* cells on average. This may be due to the more severe growth defects we observed in *est2Δ rad52Δ* compared to *est2Δ* cells. The rates of telomere shortening at each individual chromosome end were very similar in both *est2Δ* and *est2Δ rad52Δ*. Our quantitative single telomere data support previous analysis (Marcand et al. 1999) that telomere shortening rates are similar across chromosome ends.

**Figure 6.**
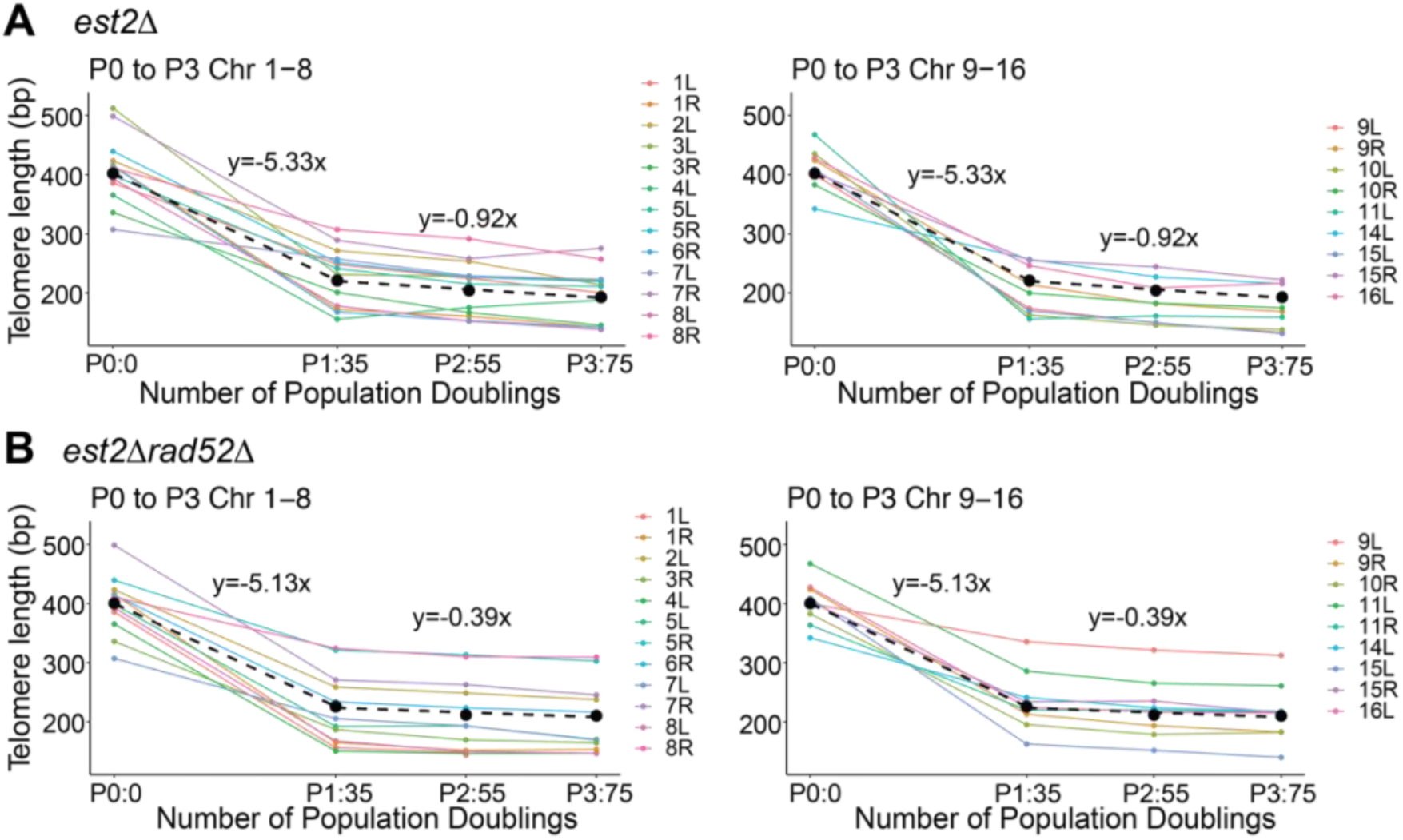
Telomere shortening rate in the absence of telomerase. A) The progression of mean telomere length at individual chromosome ends in *est2Δ* cells over 3 passages. Chromosomes 1-8 and 9-16 are shown on separate graphs for clarity. The initial telomere length, P0, is estimated from the wildtype parental strain for each chromosome, and so it is not the exact genotype (See results). The line of best fit to the average of the slopes is shown as a black dotted line. The slopes for P0 to P1 and P1 to P3 are plotted separately because the two slopes differ and each slope is reported above the line. B) The rate of telomere shortening for the *est2Δ rad52Δ* strain was measures as described above in A.

To determine the minimum telomere length in telomerase-null cells, we examined telomeres at early passages before survivor events were predicted to occur (Lundblad and Blackburn 1993). We calculated the bottom 99^th^ length percentile at each chromosome end and identified critically short telomeres in both the *est2Δ* and *est2Δ rad52Δ* ranging from 70 to 80bp. The shortest three values are highlighted in each genotype at each passage (Table 1). This minimal telomere length is also consistent with the minimal length we found in telomerase-positive wildtype and *tel1Δ* cells of ~75 bp. (Table 1). Previous work has suggested that telomeres shorter than 100 bp could be considered “critically short” (Abdallah et al. 2009). Our sequence analysis now more precisely identified a minimum of ~75 bp at individual chromosome ends.

**Table 1.** Minimum telomere length is approximately 75bp. The bottom 99^th^ percentile of telomere length at the three shortest chromosome ends per genotype at each passage in *est2Δ, est2Δ rad52Δ,* as well as for *wildtype* and *tel1Δ* are shown.

| Table 1. |  |  |  |  |  |  |  |
| --- | --- | --- | --- | --- | --- | --- | --- |
| WT |  | est2Δ P1 |  | est2Δ P2 |  | est2Δ P3 |  |
| Bottom 99% of telomere length |  |  |  |  |  |  |  |
| 13L | 77.25 | 10L | 93.2 | 10L | 79 | 10L | 77.4 |
| 7R | 96.06 | 11L | 70.4 | 11L | 71.6 | 11L | 69.4 |
| 8R | 96.19 | 2R | 72 | 2R | 75.6 | 8L | 85 |
| tel1Δ |  | est2rad52Δ P1 |  | est2rad52Δ P2 |  | est2rad52Δ P3 |  |
| Bottom 99% of telomere length |  |  |  |  |  |  |  |
| 13L | 69.3 | 10L | 86.45 | 10L | 90.4 | 10L | 76.75 |
| 4R | 73.4 | 1L | 89.6 | 1L | 81.7 | 1L | 71.9 |
| 8R | 69.4 | 3L | 171 | 2R | 85 | 2R | 71.95 |

## DISCUSSION

Here we show that nanopore sequencing can enable single molecule telomere length measurement in yeast. This novel method provides increased resolution over Southern blotting, with exquisite single molecule sensitivity and the ability to assign telomere length measurement to individual chromosome ends. Leveraging this, we were able to uncover new aspects of telomere biology. This measurement can be done on a bench top with a relatively low capital investment; even a smaller, less expensive Flongle flowcell provides a suitable number of independent reads for bulk telomere length measurement. With further improvements and the use targeted enrichment (Gilpatrick et al. 2020; Kovaka et al. 2021), the cost and yield of this method can be even further improved.

We were surprised to see apparent chromosome end specific telomere length differences preserved across multiple independent clones with different passaging histories. Previous work suggested that significant clonal changes occur after only 30 population doublings (Shampay and Blackburn 1988). If these chromosome end-specific differences are observed beyond our sample size here, then understanding how they are established and maintained would be a fascinating new area of telomere biology to explore.

The stability of the end specific lengths is surprising. While we did observe a few instances of clonal variation the majority of chromosome specific ends remained stable and did not revert to a mean length as predicted by models for telomere length maintenance. The stability of the chromosome specific differences implies that telomerase specifically elongating the shortest telomere (Teixeira et al. 2004) may only occur when there is a catastrophically short telomere. For example, we compare telomere 13L that is maintained around a mean of 241bp to telomere 7R that is maintained around 434bp (Figure 3A); if the shortest telomeres are elongated, we would expect the mean length of 13L to have increased after 120 population doublings. Perhaps, with even more population doublings, the end specific differences may revert to the grand population mean length for all ends, but our data indicate that if this occurs it is much slower than would be expected from the frequent elongation of short telomeres previously seen (Teixeira et al. 2004).

If the generation of end specific length regulation is not due to clonal heterogeneity, there are a number of models that might be explored to understand this new biology. The effects might be due to subtelomeric coding regions, subtelomeric repeats influence on telomere length, or perhaps due to specific chromatin modifications. The hint that there may be concordance between telomeres on the same chromosome (Figure 3A/C) suggests that perhaps chromosome territories within the nucleus (Noma 2017) have an effect on length regulation. Alternatively topologically associated domains (TADs) or localization to the nuclear periphery may play a role in chromosome specific telomere length differences. Nanopore sequencing of telomeres can provide the individual chromosome end resolution to further interrogate these interesting new questions.

We also found a small effect of Y’ subtelomeric elements on telomere length regulation by *tel1Δ* and *rif1Δ* mutants, indicating some role for subtelomeric sequences in length regulation. This effect was not as large as that reported by Craven and Petes (Craven and Petes 1999), but the presence of the Y’ element did appear to remove the *rif1Δ* elongated telomere phenotype in *tel1Δ rif1Δ* double mutants. This small effect is of interest, as it suggests a potential additional role of Rif1 protein on telomere length. Rif1 can exert telomere specific length regulation when it is tethered at a unique telomere (Shubin et al. 2021). Perhaps there is another binding site for Rif1 or other telomere length regulators in the X element subtelomere, and the addition of a long 5kb Y’ element moves this site too far from the telomere to exert any effect on length regulation. The detection of these subtle length differences is necessary to interpret the effects of molecular mechanisms of subtelomeric elements and other structures on telomere length regulation.

In the absence of telomerase, all telomeres shortened at a similar rate as predicted by the end replication problem. A shortening rate of about 5 bp per generation is similar to what has been seen previously with other methods. The shortest telomeres in telomerase null cells, as well as in wildtype and *tel1Δ* cells, was around 75bp. This is shorter than previously suggested (Abdallah et al. 2009), and allows specific hypotheses to be made for what might cause loss of telomere function. For instance, Rap1 binds telomeric DNA approximately every 20bp (Gilson et al. 1993; Wahlin and Cohn 2000), so telomere dysfunction might be due to loss of Rap1 binding, or perhaps loss of the terminal nucleosome binding. However, a minimal length of 75 bp should still allow Ku’s end binding function at the telomere (Gravel et al. 1998; Chen et al. 2018). Perhaps a loss of Rap1, nucleosome binding, or other telomeric binding factors in the presence of Ku allows for signaling the DNA damage response. Telomere length measurement by nanopore sequencing will allow the telomere field to study new questions and revisit past unanswered questions in telomere biology.

## METHODS

### Biological Resources

Yeast strains and plasmid vectors are listed in (Strain sheet) and are available upon request.

### Molecular Cloning

Yeast molecular cloning, culturing, sporulation, and transformation were performed as previously described in (Green and Sambrook 2012; Keener et al. 2019; Shubin et al. 2021).

The UE1 (previous named 1L 5XUAS landing pad) and UE11 (previously named 11R 5XUAS landing pad) constructs were designed *in silico* using SnapGene software and constructed as described in (Shubin et al. 2021). UE11 was generated in the same manner as UE1, but with homology at chromosome arm 11R, rather than chromosome arm 1L. The subtelomere was removed in these strains using homology to the unique gene region directly upstream of the telomere and ensuring recombination with a digested product with a 3’ overhang that is endogenously elongated by telomerase.

### Yeast strains culturing and transformation

CVy61 (W303-0 WT haploids), UE1 (previously CALy117, *1L::leu2*), UE11 (previously CALy119, *11R::leu2*). UE1 *rif1Δ* (previously CALy202, *1L::leu2, rif1Δ::kanMX*), UE11 *rif1Δ* (previously CALy204, *11R::leu2, rif1Δ::kanMX*), and UE1 *tel1Δ* (previously CALy472, *1L::leu2, tel1Δ:: hphMX4*) strains were generated as previously described in (Shubin et al. 2021). These strains were re-named UE1 and UE11 to highlight the function of a unique end, rather than a 5XUAS landing pad, that was important to this study. A list of all yeast strains and genotypes is presented in Supplemental Table 4.

The UE1 *tel1Δrif1Δ* was generated by transformation. Briefly, transformation was performed by treating 50mL of logarithmically growing UE1 *rif1Δ* cells with 0.1M lithium acetate (L6883; Sigma) and adding 1ug of linear DNA to replace *TEL1* with *tel1Δ::hphMX4* cassette for one-step integration. Cells were incubated at 30C for 10 minutes before the addition of 0.5 ml 40% polyethylene glycol (PEG4000, P4338; Sigma),0.1M lithium acetate. Cells were then incubated for 30min at 30C followed by a 45min heat shock at 42C. Cells were plated after a water wash onto YPD + HygromycinB, (Corning, at 200ug/ml). Integration at the *TEL1* locus was confirmed by junction polymerase chain reaction (PCR). A list of primers and plasmids used in this study is presented in Supplemental Tables 5 and 6, respectively.

To perform the W303-0,1,2,3 experiment, CVy61 was streaked to single colony on YPD media. One colony was selected (W303-0) and grown in 5mL YPD to 0.5 OD. This culture was split into two subcultures: 1) growth to 1.0 OD in 100 mL YPD broth for High Molecular Weight genomic DNA extraction (see below) and called W303-0 for nanopore sequencing and 2) growth in 10mL YPD broth to saturation for 24 hours and passaged as 5uL of saturated culture in 10mL fresh YPD. This was repeated a total of 5 times before cells were plated onto YPD media. Three independent colonies were selected (W303-1, W303-2, and W303-3) to inoculate three cultures of 100 mL YPD broth for DNA extraction as described for subculture #1. We estimate 120 population doublings between W303-0 and W303-1,2,3. See Supplemental Figure S6A for illustration.

To perform the UE1 and UE11 experiments in wildtype and *rif1 Δ* backgrounds, frozen stocks of UE1 wildtype, UE11 wildtype, UE1 *rif1Δ,* and UE11 *rif1Δ* were streaked onto YPD media to single colony. One colony each was selected from UE1 wildtype, UE11 wildtype, UE1 *rif1Δ,* and UE11 *rif1Δ,* grown to 0.5 OD in 5mL YPD broth, followed by growth to 1 OD in 100 mL YPD for DNA extraction and nanopore sequencing as described above. An additional colony was selected from UE1 wildtype and UE1 *rif1Δ* and grown in 10mL YPD to saturation for 24 hours and passaged as 5uL of saturated culture in 10mL fresh YPD. This was repeated a total of 3 times in liquid culture before cells were grown to 0.5 OD in 5mL YPD broth, then to 1.0 OD in 100mL YPD broth for DNA extraction and sequencing of UE1-2 wildtype and *rif1Δ.* We estimate 60 population doublings between UE1-1 and UE1-2. See Supplemental Figure S6B for illustration.

To generate telomerase-null *est2Δ* and *est2Δ rad52Δ* cells, the heterozygous diploid strain yAY139 (IJpma and Greider 2003) was sporulated and dissected to yield sister spores for analysis.

Since these mutants will senesce with passaging, we attempted to keep cell division to a minimum, allowing telomeres to shorten, but not form survivors (Chen et al. 2001). For both *est2Δ* and *est2Δ rad52Δ,* cells were taken directly from the original dissection plate, and grown to saturation in 10mL cultures of YPD broth. These mutant cultures were: 1) grown to 1 OD in 100 mL for DNA extraction and sequencing, called passage 1 (P1) or 2) diluted to 10^5^ cells/mL in YPD broth every 24 hours before being grown to 1 OD in 100 mL for DNA extraction and sequencing. Dilution and growth for these mutants was continued for two more rounds, collecting cells after each passage called P2 and P3. See Supplemental Figure S10 for illustration. A wildtype haploid spore was also selected from the original dissection plate of yAY139, grown to saturation in 10mL YPD, and then grown to 1 OD in 100mL YPD for DNA extraction and sequencing. This sample was called passage 0 (P0).

### High molecular weight genomic DNA extraction

We modified a DNA isolation protocol (Denis 2018) to collect high molecular weight (HMW) genomic DNA from yeast. Cultures were grown at 30C to 1.0 OD in 100 mL YPD broth, collected in a Sorvall Legend XTR centrifuge with swinging bucket rotor for 5min at 1500xg and washed once with 20mL TE buffer (10mM Tris, 1mM EDTA, pH8.0).

To generate spheroplasts, cell pellets in a 50ml conical centrifuge tube (Falcon) were resuspended in 2mL 1M sorbitol with 0.5 mL of 2.5 mg/mL Zymolyase 100T (Amsbio, 120493-1) and shaken gently (75rpm) at 30C for 1hour. Spheroplasts were collected at 300xg for 4min in the swinging bucket centrifuge and the supernatant was carefully discarded.

Pelleted spheroplasts were lysed by resuspension in 1.7 mL lysis buffer (100mM Tris, pH7.5, 100mM EDTA, pH8.0, 500mM NaCl, 1% PVP40) and 250μL10% SDS. After inverting the tube gently10 times, 20μL of 100mg/mL RNaseA (Sigma, R6513) was added and the mixture was shaken at 75 rpm at 37C for 30min. To digest protein, 25μL of 20mg/mL Proteinase K (Invitrogen 25530-015)) was added, and the mixture was incubated for 2hr at 50C with occasional inversion until there was a uniform cloudy suspension.

To precipitate and remove protein, 5mL TE and 2.5 mL 5M potassium acetate, pH7.5 were added to the suspension and the tube was inverted 10 times to mix, then placed on ice for 5min. The tube was inverted again and placed on ice for an additional5min before centrifugation at 4C for 15min at 4000xg in the swinging bucket centrifuge. The supernatant was decanted into a clean tube and the centrifugation was repeated. The supernatant was again decanted into a clean tube.

DNA was precipitated by adding an equal volume of 100% isopropanol and swirling the tube until the DNA strands are observed. Precipitated DNA was collected from the bottom of the tube using a p1000 wide bore pipette tip and placed in a clean Eppendorf microcentrifuge tube. The DNA was pelleted in a minicentrifuge for 15sec and the supernatant carefully decanted. 500 uL 75% ethanol was added to rinse the DNA pellet. After decanting the wash, the residual ethanol was removed with a pipette tip. The DNA pellet was resuspended in 50-200 uL of EB buffer (Qiagen, 10mM Tris-Cl, pH8.5), depending on the size of the pellet. The DNA was allowed to resuspend at 4C on an inverting rotator overnight. The final concentration of resuspended DNA was measured on a Qubit 3.0 Fluorometer (Thermo Fisher Scientific).

### DNA molecular end tagging

To tag the molecular end, HMW genomic DNA (gDNA, see above) was first A-tailed using terminal transferase (NEB M0315) in a reaction by incubating 2ug of gDNA, 1X terminal transferase buffer, 1X CoCl_2_, 5mM dATP, and 20U of terminal transferase enzyme for 1hr at 37C, followed by heating at 70C for 10min to stop the reaction.

Five primers were designed to make a mixture of TeloTags (see Primer Table).

To the A-tailed gDNA reaction, the following were added to complete the TeloTag addition: 1x ThermoPol Reaction Buffer Pack (NEB B9004S), 2.5mM dNTP mix, 1mM ATP, 2.5mM TeloTag primer mix, and 4units of Sulfolobus DNA Polymerase IV(NEB M0327S). The TeloTag reaction was incubated in a Veriti 96 well Thermal Cycler (Applied Biosystems) for 1min at 56C and 10min at 72C. 400 units of T4 ligase (NEB M0202) was then added to the reaction, and allowed to continue incubating at 12C for 20min. Ampure XP beads (Beckman Coulter) were used to purify the tagged gDNA from the reaction mixture.

Size selection of the tagged gDNA was performed using the Short Read Eliminator XS kit (Circulomics, SS-100-121-01), which retains DNA molecules greater than 5kb. Once this step is completed, the tagged gDNA is ready for nanopore library preparation (see below).

### Southern blotting and densitometry

HMW genomic DNA was extracted and quantitated as described above. Telomere length analysis by Southern Blot was performed on the same strains used for nanopore sequencing according to protocol described in (Kaizer et al. 2015). Briefly, 250 ng of gDNA was digested with PvuII and XhoI for 2-4 hours at 37C and resolved on a 1% agarose gel overnight. Two concentrations of 2-log ladders (NEB) were included as reference for analysis, 10ng for single telomere assays and for bulk telomere assays. After transfer, the membrane was hybridized with ^32^P radiolabeled 2-log ladder and PCR fragments unique to either 1L, 11R (a purified PCR product of the LEU2 gene and CYC1 terminator as described in Shubin, et al.), Y’ subtelomeric elements as described in Kaizer, et al. or to CEN4 (Kaizer et al. 2015; Laterreur et al. 2018; Shubin et al. 2021). After washing, the membrane was exposed on a Storage Phosphor screen (GE Healthcare) for 1 day for bulk assays and 4-5 days for single telomere assays. Southern images were captured on a STORM using ImageQuant (GE Healthcare). ImageQuant .gel files were downloaded into Adobe PhotoShop CS6 and saved as .tif files.

Southern blot densitometry was measured to allow for comparison to Nanopore data as it shows both the normal distribution and spread of telomere length for each sample. However, these plots are often inconsistent and are not a direct measure of telomere length as they are generated from the Southern blot image. To somewhat counter this, we measured densitometry using two programs. We first used ImageQuant to generate a distribution fully across each sample lane to be overlaid with Nanopore telomere fragment length data, so that all molecular markers and the resolution cut-off would be visible (Fig. 2C, D, F, G). In ImageQuant, we used the 2-log ladder and the CEN4 band at 1.4kb as molecular markers to determine a linear range and the resulting .csv files were loaded into RStudio to generate telomere length plots. Second, PhotoShop .tif files were inverted and loaded into WALTER ScanToItensity and IntensityAnalyzer to remove background and generate intensity profiles of the telomere signal alone (Lycka et al. 2021). The reported summary statistics were loaded into RStudio to produce boxplots of telomere length.

### Nanopore library preparation and sequencing

Library preparation of 1μg input DNA with a molecular tag was performed using the Native barcoding genomic DNA kit (Oxford Nanopore Technologies EXP-NBD104 and SQK-LSK109). Samples were run on a MinION flow cell (v.9.4.1) or Flongle flow cell (v.9.4.1 pore) using the MK1B or GridION for 24 to 72 hours, and were operated using the MinKNOW software (v.19.2.2). Only reads with barcodes on both ends were selected to pass. The read counts and characteristics are shown in Supplemental Tables S1, S2, S3. We note that this WGS sequencing from a MinIon flow cell provides enough coverage (~50-200 reads per telomere) to analyze specific single chromosome ends, but Flongle flow cells produce only ~10 reads at a single chromosome end, which at present are not enough to analyze specific single chromosome ends. In the future, enrichment methods could be used to increase the yield of telomere-specific reads, and a Flongle flow cell may be sufficient to analyze specific chromosome ends (Gilpatrick et al. 2020). We note future studies can use this nanopore method to correct the subtelomere assemblies in various strains, however this was out of the scope for this study.

### Nanopore analysis, alignment and telomere length calculation

Base calling to generate FASTQ sequencing files from the electrical signal data (FAST5 files) was performed with GUPPY (v.4.2.3). FASTQ reads containing telomere sequence were selected using TideHunter (v1.4.4) (Gao et al. 2019). Reads less than 5kb were filtered out using awk. Reads containing the TeloTag were selected using seqkit (v0.16.1) (Shen et al. 2016). Nanopore adapters, barcodes, and the TeloTag were removed from reads using PoreChop (v0.2.4) (Wick 2018). FASTQ sequence reads were aligned to a custom reference genome modified from sacCer3(Foury et al. 1998) (where 2kb of synthetic telomere repeat sequence was added to each chromosome end) using minimap2 (v2.18) (Li 2018). Custom references for WT W303-0, UE1, UE11, and S288C are available below. Secondary and supplementary reads were filtered out using samtools (v1.12), so that only high confidence alignment primary reads were selected. Bedtools (v2.30.0) was used to generate files with the start and end chromosomal position of each read (Quinlan and Hall 2010). These bedfiles and a bedfile with the start position of each telomere end (available for WTW303, UE1, and UE11 references at below) were read into a custom script in RStudio (available below) to calculate telomere length for each read and generate plots using the R package ggplot2 (v3.3.3) (Wichman et al. 2016). Statistics were calculated using the R package rstatix (v0.7.0) and ANOM (v0.2)(Pallmann 2017; Kassambara 2021).

## Code Availability

The reference genomes, telomere start position bedfiles, and code for all the above analysis is hosted on GitHub (https://github.com/timplab/Yeast-Telomere-Nanopore.git).

## DATA ACCESS

Sequencing data for this study can be retrieved from the SRA, under the BioProject ID PRJNA730563.

## COMPETEING INTEREST STATEMENT

W.T. has two patents (8,748,091 and 8,394,584) licensed to Oxford Nanopore Technologies.

## ACKNOWLEDGEMENTS

This study was supported by the NIH R35CA209974 (to CWG) and NIGMS T32 GM007445 (to the BCMB graduate training program).

We acknowledge Carla Connelly for strain generation and Southern blotting. We thank Dr. Brendan Cormack, Dr. Rebecca Keener, and Dr. Calla Shubin for thoughtful discussions about experimental design and conclusions. We would like to thank Carla Connelly, Dr. Brendan Cormack, Dr. Rebecca Keener, Dr. Paul Hook, Dr. Akshi Jasani, and Thea Egelhofer for critical reading and editing of the manuscript.

