## Supplemental Material for "Chromosome specific telomere lengths and the minimal functional telomere revealed by nanopore sequencing"

#### Table of contents


##### Supplemental data:

**Supplemental Figure S1 –** Subtelomere structure for UE1 and UE11

**Supplemental Figure S2** – Telomere fragment length measured by Nanopore sequencing accurately reflects length measurements by Southern blot in UE11

**Supplemental Figure S3** – Quantitative analysis of nanopore sequencing compared to Southern blot for UE1 Y' telomeres and unique End 1L

**Supplemental Figure S4** – Telomere fragment length measured by Nanopore sequencing is a normal distribution and has a good correlation to measurements by Southern blot

**Supplemental Figure S5** – Nanopore telomere length measurement works reproducibly and on a smaller flow cell

**Supplemental Figure S6** – UE1 and UE11 lineage and clonal selection

**Supplemental Figure S7 –** Analysis of the means (ANOM) multiple contrast test of each telomere against the grand mean of all telomere length profiles

**Supplemental Figure S8 –** W303 clonal selection, passaging, and telomere length correlations

**Supplemental Figure S9:** *rif1Δ* individual chromosome end length profiles are not gaussian and do not correlate across replicates

**Supplemental Figure S10 –** Schematic of clone selection after sporulation and tetrad dissection of an EST2::est2Δ RAD52::rad52Δ diploid

##### Supplemental Tables:

**Supplemental Table 1** – Read count for whole genome collection, telomere-specific, and filtered reads for telomere length analysis for 1L and 11R strains in Fig. 2

**Supplemental Table 2** – Read count for whole genome collection, telomere-specific, and filtered reads for telomere length analysis 11R strains run on a MinIon versus Flongle in Supplemental Figure 5

**Supplemental Table 3** – Read count (N) for telomere-specific reads, mean, median, and standard deviation values reported for telomere length analysis of all WT, est2Δ, and est2Δ rad52Δ cells over ~75 population doublings (P1-P3)

**Supplemental Table 4** – Yeast strains used in this study

**Supplemental Table 5** – Primers used in this study

**Supplemental Table 6** – Plasmids used in this study

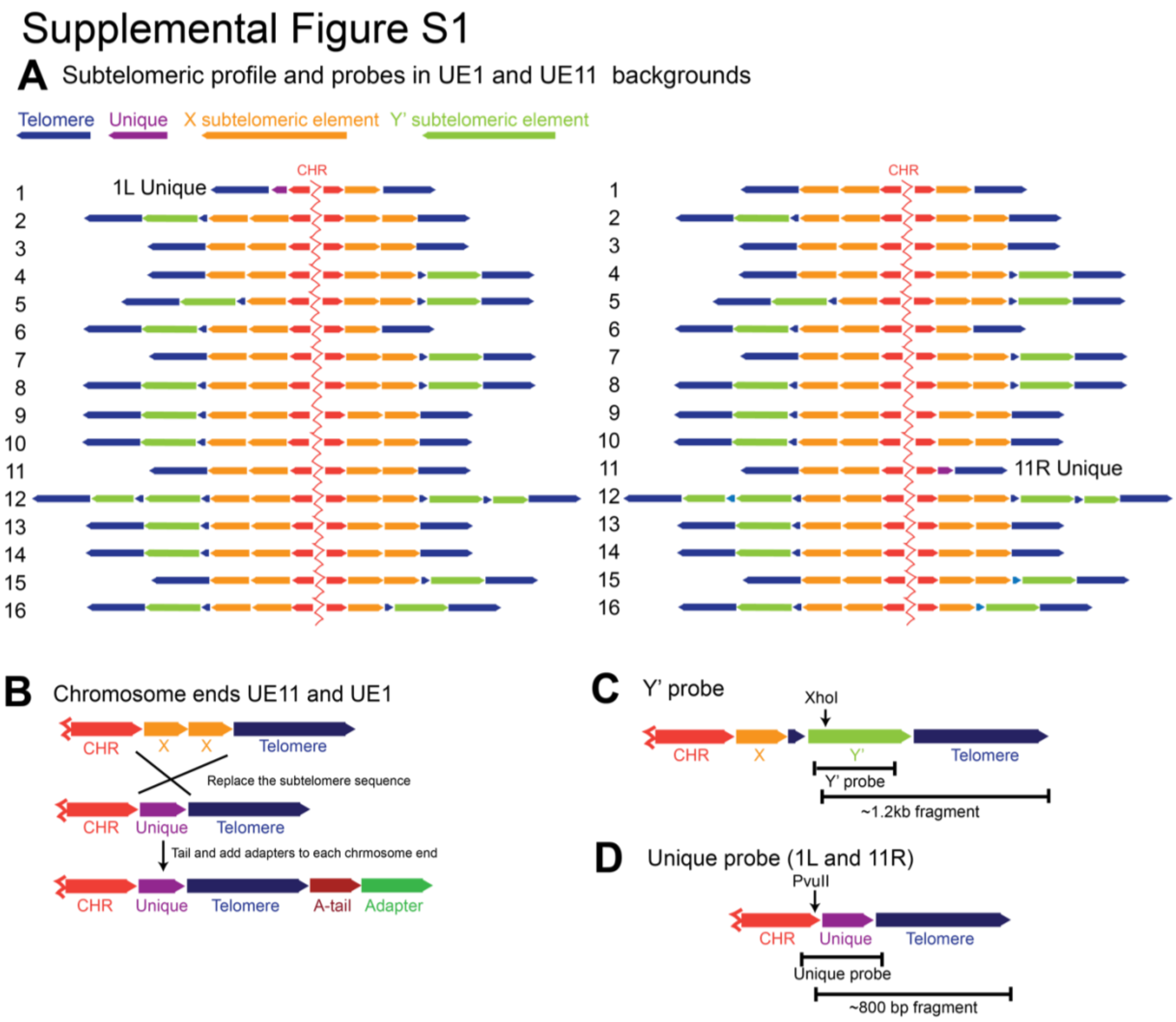


**Supplemental Figure S1. Subtelomere structure for UE1 and UE11**. A) Schematic of the subtelomeric sequence structure for strains with unique chromosome end generation by replacing the subtelomere at 1L and 11L. All yeast chromosomes have an X element in the subtelomere and some chromosomes also have one or more Y' elements. Between the X and the Y' elements and between multiple Y' elements there is ~100 bp of $G_{1\text{-}3}T$ telomere repeats (Louis et al. 1994; Louis 1995). (Louis et al. 1994; Louis 1995). Y' elements are relics of retrotransposons that now move by recombination (Louis and Haber 1992). Thus different yeast strains have very different subtelomere structure, as shown. B) Schematic of how strains with unique ends at 1L and 11R were generated by transformation. C) The expected size of the bulk Y' containing telomeres after cutting with XhoI for and hybridization with Y' probe. D) The expected size of the unique telomere ends after cutting with PvuII and hybridization with Unique telomere probe.

#### Supplemental Figure S2

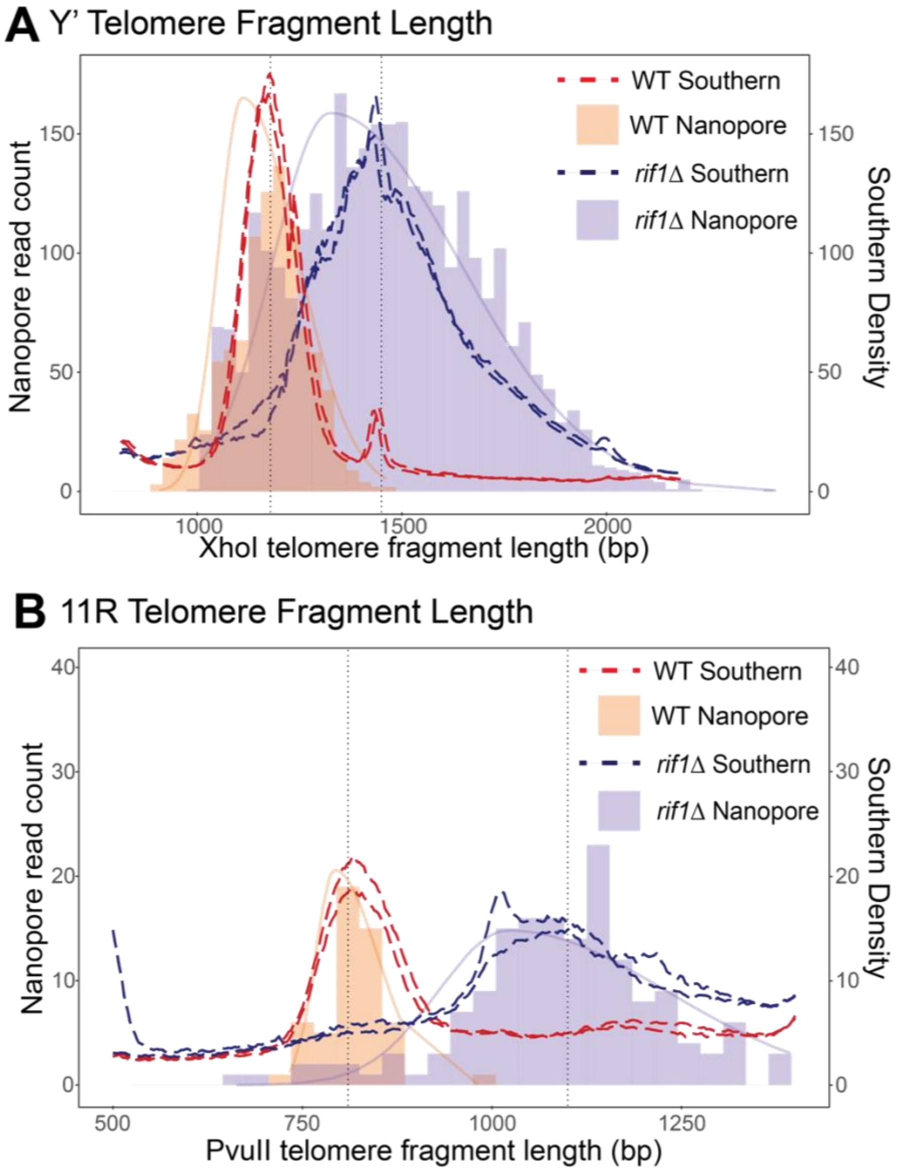


**Supplemental Figure S2. Telomere fragment length measured by nanopore sequencing accurately reflects length measurements by Southern blot in UE11.** A) Comparison of densitometry (dotted lines) of Southern shown in (Fig. 2A) and nanopore sequencing (bars). The light orange bars represent nanopore read counts for WT cells, the light purple bars represent nanopore read counts for *rif1Δ* cells. C) Comparison of densitometry (dotted lines) of Southern and nanopore sequencing (bars) of 11R telomeres. The light orange bars represent nanopore read counts for WT cells, the light purple bars represent nanopore read counts for *rif1Δ* cells.

#### Supplemental Figure S3

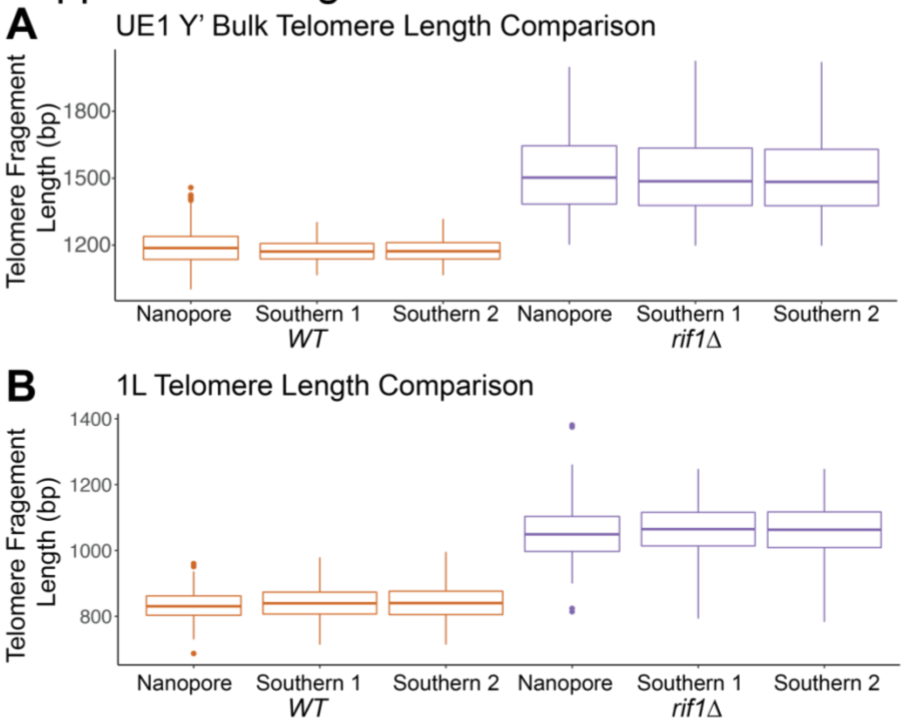


**Supplemental Figure S3. Quantitative analysis of nanopore sequencing compared to Southern blot for UE1 Y' telomeres and unique End 1L.** Densitometry by WALTER analysis of terminal restriction fragments (TRF) length (Lycka et al. 2021) was performed on the A) UE1 Y' bulk telomeres and B) UE1 1L telomere for WT and *rif1Δ* samples in duplicate to generate intensity profiles and summary statistics of telomere length (mean, median, and standard deviation, shown as WT Southern 1 and 2 and *rif1Δ* Southern 1 and 2). These were plotted against nanopore fragment analysis (*rif1Δ* Nanopore and WT Nanopore).

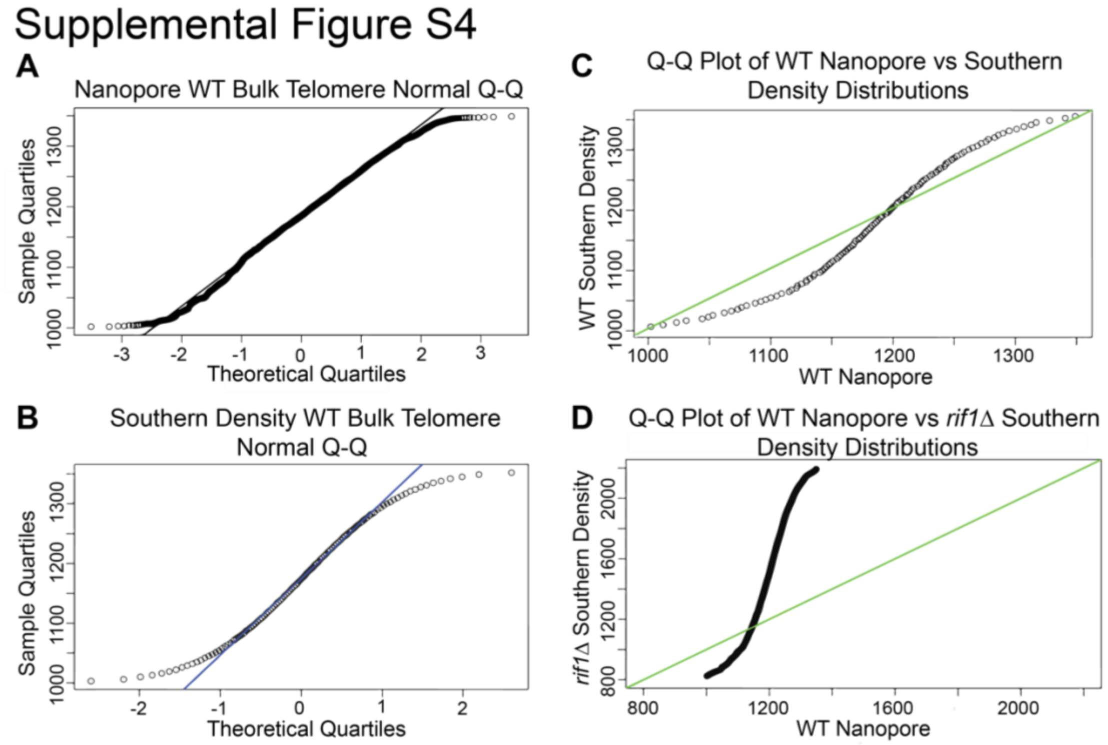


**Supplemental Figure S4. Telomere fragment length measured by Nanopore sequencing is a normal distribution and has a good correlation to measurements by Southern blot .** A) Normal Q-Q plot for Nanopore WT Y' telomere fragment measurement plotted against a normal distribution, demonstrating agreement into the 2nd quartile. B) Normal Q-Q plot for Southern blot densitometry WT Y' telomere fragment measurement plotted against a normal distribution, demonstrating agreement into the 1st quartile. The increased thin tails present on the Normal Southern Q-Q plot versus the Normal nanopore Q-Q plot suggest an increased dynamic range of measurements from the nanopore. data. C) Q-Q plot for correlation between nanopore and Southern blot densitometry for WT Y' telomere fragment measurements. The linear response of this plot shows the similarity between the measurement methods. D) Control Q-Q plot of a *rif1Δ* Southern dataset and WT Nanopore dataset that shows a clear deviation from the expected linear relationship.

#### Supplemental Figure S5

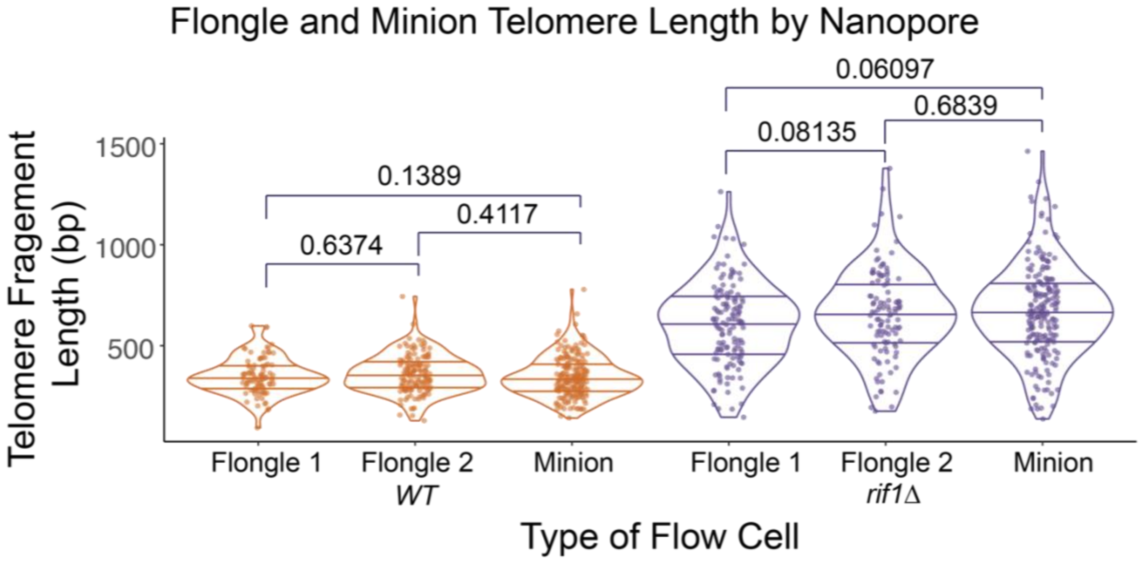


**Supplemental Figure S5. Nanopore telomere length measurement works reproducibly and on a smaller flow cell.** Telomere length comparison between MinIon and Flongle flow cells on the same WT and *rif1Δ* samples in strain UE11 background. The minion datasets were randomly down sampled to N=200 to have a comparable number of reads to the flongle datasets. A 2-sample t-test was run on each pair of samples of the same genotype. No p value was less than 0.05, indicating that there are no meaningful statistical differences between Flongle and MinIon flowcells within a genotype.

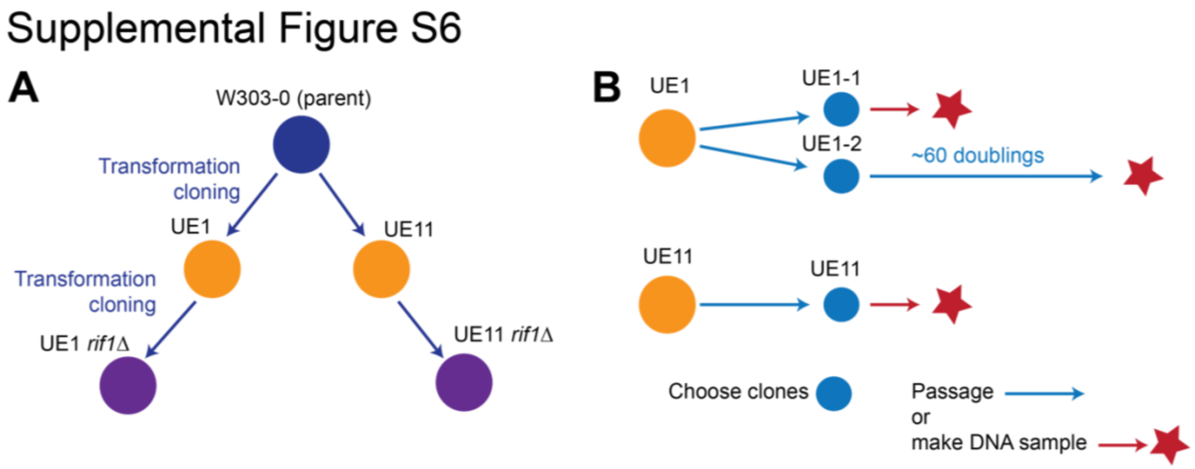


**Supplemental Figure S6**. **UE1 and UE11 lineage and clonal selection.** A) Schematic of the linage from parental W303-0 to generate UE1 and UE11 WT strains (orange) and then transformation to generate UE1 and UE11 *rif1Δ* (purple). B) Schematic of independent clone selection from daughter UE1 and UE11 strains showing additional passages before nanopore sequencing. This selection was repeated for UE1-*rif1Δ* and UE11-*rif1Δ* strains.

Supplemental Figure S7

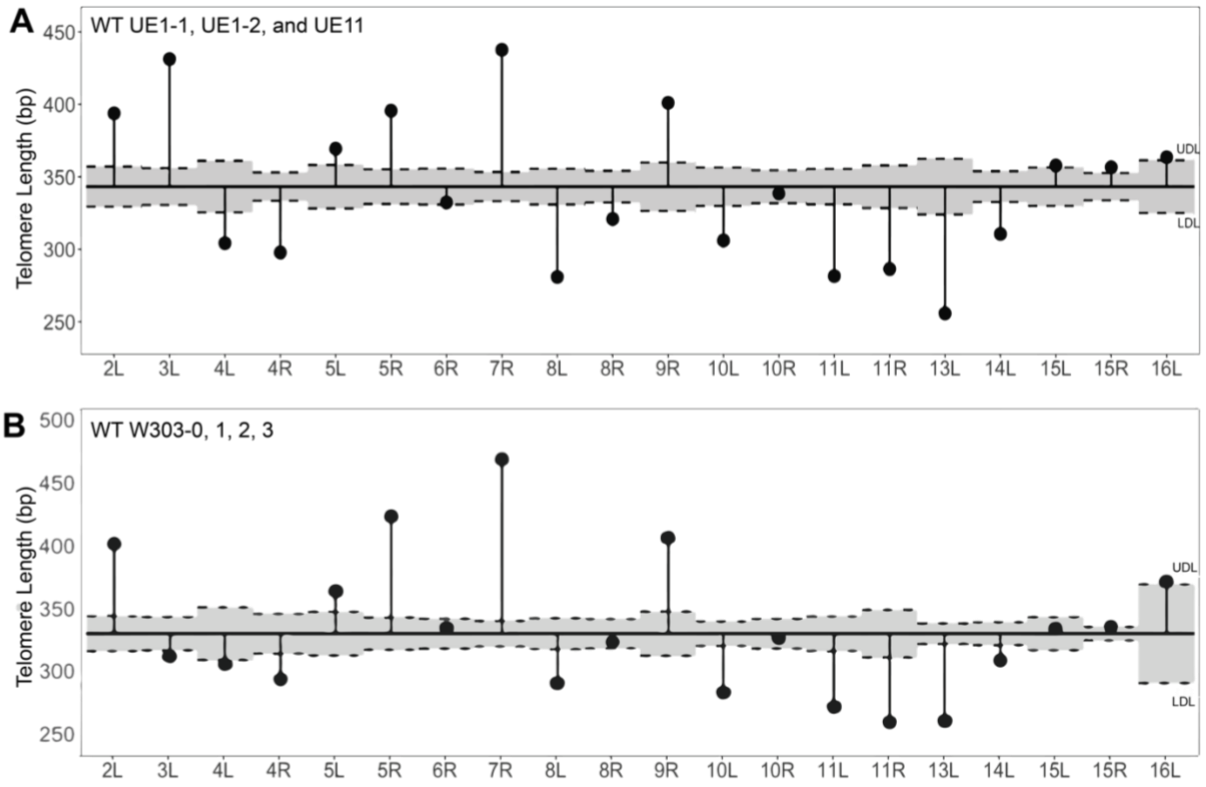


**Supplemental Figure 7. Analysis of the means (ANOM) multiple contrast test of each telomere against the grand mean of all telomere length profiles.** WT ANOM analysis of A) UE1-1, UE1-2, and UE11 and B) W303-0, 1, 2, 3 telomere length distributions against the grand mean telomere length (343.3 bp). P values were adjusted using the Bonferroni method, and chromosome ends with length profiles reaching outside of the shaded grey region between the upper decision limit (UDL) and lower decision limit (LDL) are considered significantly different from the grand mean.

#### Supplemental Figure S8

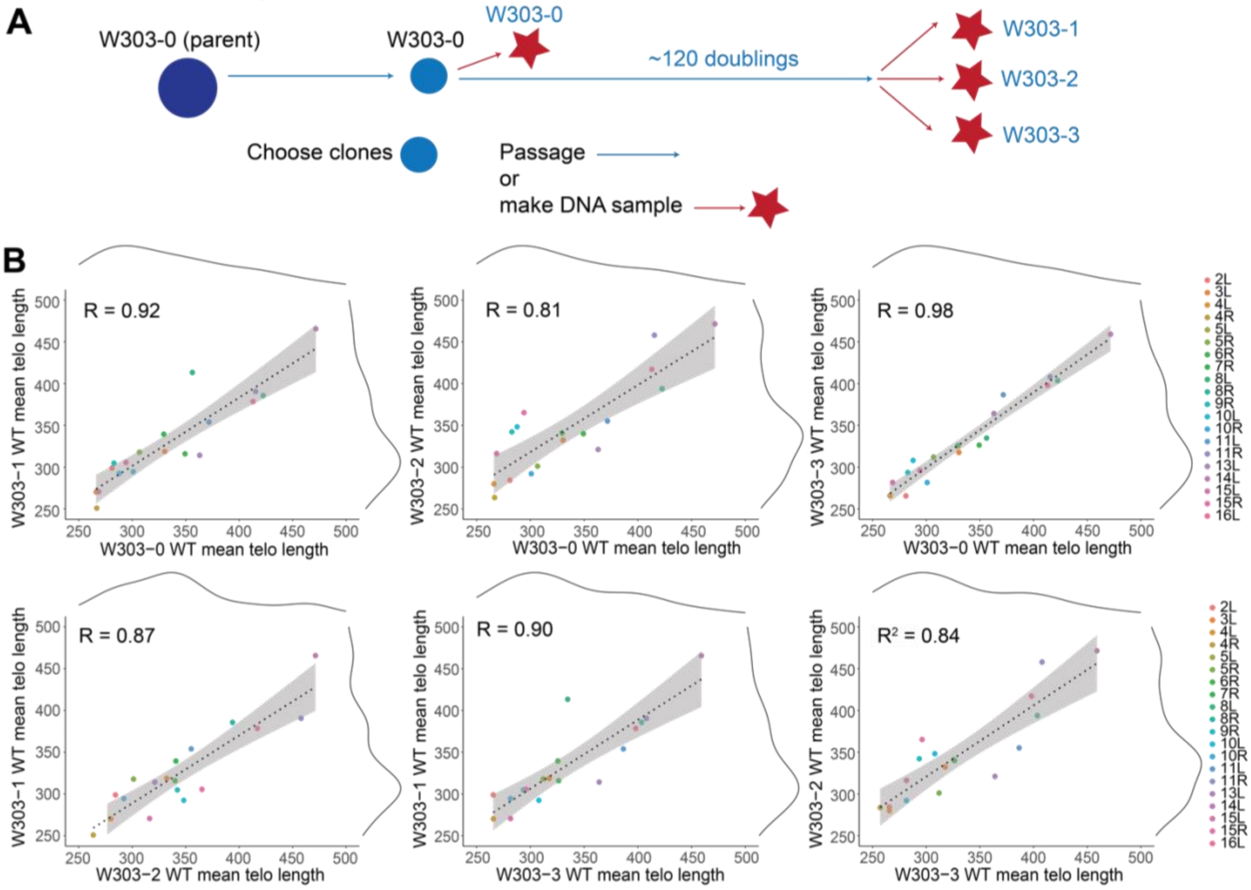


**Supplemental Figure S8. W303 clonal selection, passaging, and telomere length correlations**. A) Schematic of independent clone selection from the parent W303 WT strain and additional passages before nanopore sequencing. B) Pairwise correlations of mean telomere length across each of the four replicates for W303 WT cells. A linear model was fit to the data (dotted line) to calculate the correlation coefficient, reported as the $R^2$ value. Marginal density of each dataset is plotted as solid black lines across the top and right of the plots.

#### Supplemental Figure S9

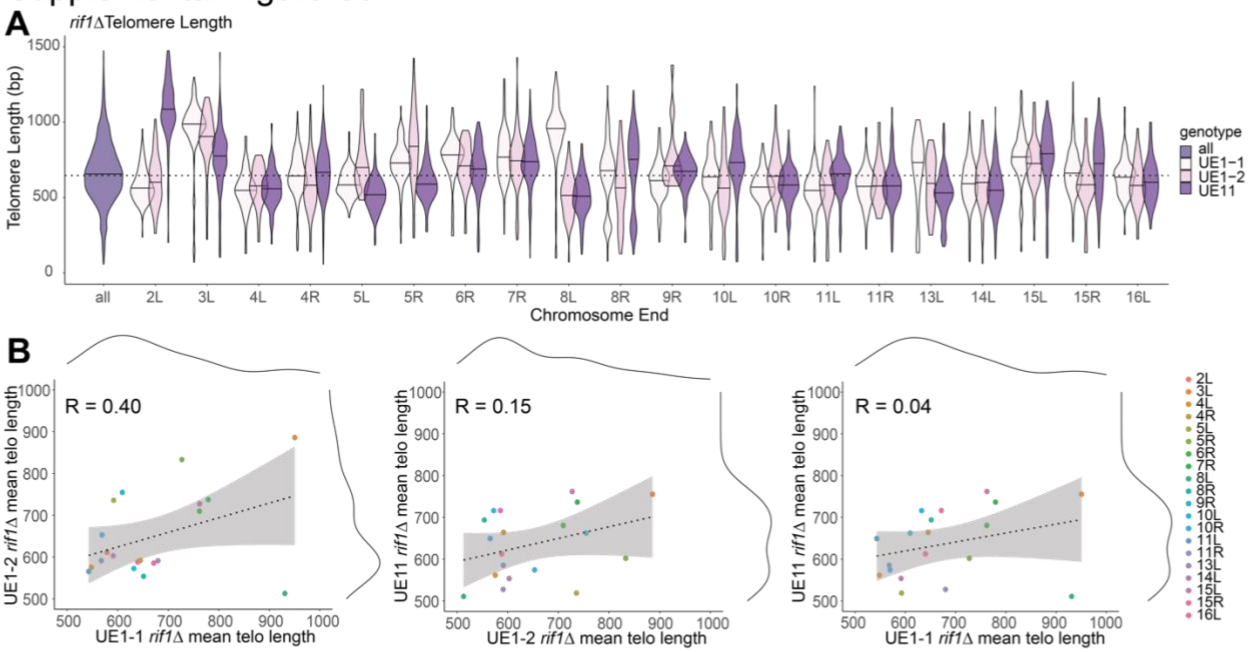


**Supplemental Figure S9: *rif1Δ* individual chromosome end length profiles are not gaussian and do not correlate across replicates.** A) Bulk telomere length (all) and telomere length for each chromosome end were mapped by aligning to internal unique sequence for *rif1Δ* cells. Three independent clones were sequenced two from UE1 (UE1-1 and UE1-2) and one from UE11 (see supplemental figure S6). Telomere lengths are reported as violin plots with a line at the mean. A dotted line shows the mean of the total population at 667 for *rif1Δ* cells. B) Pairwise correlations of mean telomere length across each of the four replicates for W303 WT cells. A linear model was fit to the data (dotted line) to calculate the correlation coefficient, reported as the $R^2$ value. Marginal density of each dataset is plotted as solid black lines across the top and right of the plots.

### Supplemental Figure S10

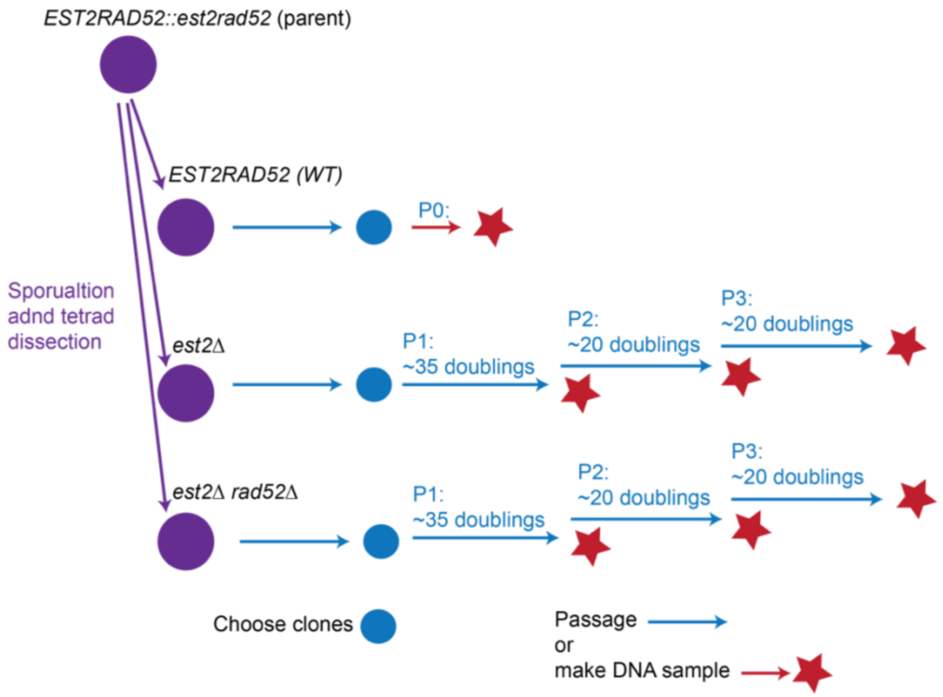


**Supplemental Figure S10. Schematic of clone selection after sporulation and tetrad dissection of an *EST2::est2Δ RAD52::rad52Δ* diploid.** WT cells were used in the analysis as Passage 0 (P0). Passage 1 (P1) is about 35 doublings, Passage 2 (P2) is 55 doublings, and Passage 3 (P3) is 75 doublings.

#### Supplemental Table S1

| Yeast Strain | Total reads (per barcode) | Gb yield (per) | N50 (kb) | Median read length (kb) | Total telomere-containing reads (%) | Primary long telomere reads for length calculation (%) |
|---|---|---|---|---|---|---|
| UE1 WT | 1.4 M | 7.34 | 13.34 | 14.16 | 6537 (0.46) | 4744 (0.34) |
| UE1 *rif1Δ* | 1.5 M | 7.87 | 13.34 | 14.16 | 9822 (0.65) | 5275 (0.35) |
| UE11 WT | 350,000 | 6.3 | 12.79 | 13.97 | 2628 (0.75) | 2056 (0.59) |
| UE11 *rif1Δ* | 760,000 | 6.92 | 12.79 | 13.97 | 9667 (1.27) | 6437 (1.27) |
| UE1 1L single | - | | - | - | - | 142 |
| UE1 1L single *rif1Δ* | - | | - | - | - | 144 |
| UE11 11R single | - | | - | - | - | 161 |
| UE11 11R single *rif1Δ* | - | | - | - | - | 258 |

**Supplemental Table S1. Read count for whole genome sequencing, telomere-specific, and filtered reads for telomere length analysis for 1L and 11R strains shown in Fig. 2.** Telomere sequence is calculated to be 0.08% of the yeast genome. The last 5kb of each chromosome is calculated to be 1.33% of the yeast genome.

#### Supplemental Table S2

| Yeast Strain | Total reads | Total telomere reads | Unique long telomere reads for length calculation | Mean measured telomere length | Med measured telomere length | Std measured telomere length |
|---|---|---|---|---|---|---|
| UE11 WT Minion | 350,000 | 2628 | 2056 | 342 | 331 | 95 |
| UE11 *rif1Δ* Minion | 760,000 | 9667 | 6437 | 648 | 651 | 219 |
| UE11 WT Flongle 1 | 13,000 | 161 | 143 | 349 | 340 | 90 |
| UE11 *rif1Δ* Flongle 2 | 15,000 | 258 | 211 | 628 | 634 | 205 |
| UE11 WT Flongle 1 | 36,000 | 265 | 201 | 355 | 355 | 93 |
| UE11 *rif1Δ* Flongle 2 | 31,000 | 227 | 152 | 657 | 665 | 224 |

**Supplemental Table S2.** Read count for whole genome collection, telomere-specific, and filtered reads for telomere length analysis 11R strains run on a minion versus flongle in Supplemental Figure 5.

#### Supplemental Table S3

| **Yeast Strain** | **Total telomere reads** | **Unique long telomere reads for length calculation** | **Mean measured telomere length** | **Median measured telomere length** | **Std measured telomere length** |
|---|---|---|---|---|---|
| WT | 3619 | 2372 | 406 | 401 | 96 |
| *est2Δ* Passage 1 | 4885 | 3405 | 239 | 237 | 79 |
| *est2Δ* Passage 1 run 2 | 4379 | 3107 | 241 | 236 | 81 |
| *est2Δ* Passage 2 | 4388 | 2931 | 229 | 217 | 83 |
| *est2Δ* Passage 3 | 3281 | 2463 | 228 | 213 | 88 |
| *est2Δ rad52Δ* Passage 1 | 563,000 | 3739 | 2813 | 247 | 74 |
| *est2Δ rad52Δ* Passage 2 | 440,000 | 3004 | 2277 | 238 | 77 |
| *est2Δ rad52Δ* Passage 3 | 190,00 | 1107 | 896 | 214 | 73 |

**Supplemental Table S3.** Read count (N) for telomere-specific reads, mean, median, and standard deviation values reported for telomere length analysis of all WT, *est2Δ, and est2Δ rad2Δ* cells over ~75 population doublings (P1-P3).

#### Supplemental Table S4

| **Strain ID** | **Genotype** | **Source** |
|---|---|---|
| CVy61 (W303-0) | *MATa ade2-1 trp1-1 ura3-1 leu2-3,112 his3-11,15 can1-100 RAD5 bar1Δ::hisG BrdU-inc::TRP1* | Viggiani and Aparicio 2006 |
| CALy11 | CVy61 *rif1Δ::kanMX* | Shubin, et al 2020 |
| UE1 | (Previously CALy117), CVy61 1L V2 landing pad | Shubin, et al 2020 |
| UE11 | (Previously CALy119), 11R V2 landing pad | Shubin, et al 2020 |
| UE1 rif1Δ | (Previously CALy202), UE1 *rif1Δ::kanMX* | Shubin, et al 2020 |
| UE11 rif1Δ | (Previously CALy204), UE11 *rif1Δ::kanMX* | Shubin, et al 2020 |
| JHUy783 | *MATa/MATα ura3Δ0/ura3Δ0 leu2Δ0/leu2Δ0 his3Δ1/his3Δ1 lys2Δ0/lys2Δ0 EST1/EST1-13myc::kanMX TEL1/tel1Δ::hphMX4* | Greider, unpublished |
| UE1 tel1Δ | (Previously CALy472) UE1 *tel1Δ::hphMX4* | Shubin, et al 2020 |
| UE1 rif1Δ tel1Δ | UE1 *rif1Δ::kanMX tel1Δ::hphMX4* | This study |
| yAY139 | *MATa/MATα ura3-52/ura3-52 lys-801/lys2-01 ade2-101/ade2-101 his3Δ200/his3Δ200 trp1Δ1/trp1Δ1 leu2Δ1/leu2Δ1 EST2/est2Δ::HIS3 RAD52/rad52Δ::TRP1 CFIII(CEN3.L)URA3 SUP11* | IJpma and Greider 2003 |

**Supplemental Table S4.** Yeast strains used in this study.

#### Supplemental Table S5

| Primer ID | 5'-3' SEQUENCE | Function | Source |
|---|---|---|---|
| OCC23 | TTTTCAGTTCTTTGTGTTTTTCCTC | A primer for YBR275C ( RIF1) | Winzeler, et al. 1999 |
| OCC25 | CACATGATATTATGAGCGTGATAGG | A primer for YBL088C (TEL1) | Winzeler, et al. 1999 |
| OCC80 | ATCTACGTCGATTTCTTTCATTTTG | D primer for YBL088C ( TEL1) | Winzeler, et al. 1999 |
| OCC84 | GATAAGGATGCCAATATTAATGACG | D primer for YBR275C ( RIF1) | Winzeler, et al. 1999 |
| CS153 | TAGCACCTGCATGAGTGCGAGCCACTACCATATTGTTTCTAACTGTGGGAATACTCAGGT | | This study |
| CS154 | CTCGAGGAAATAGGATGCCATTGTATCATAGCACCTGCATGAGTGCGAGCCACT | | This study |
| TeloTag Mix | CAGTCTACACATATTCTCTGTTTTTTTTTTTTTTTTTTTTTTTTTTTTTTTTT | | This study |
| | CAGTCTACACATATTCTCTGTTTTTTTTTTTTTTTTTTTTTTTTTTTTTTTTTTTTCAC | | This study |
| | CAGTCTACACATATTCTCTGTTTTTTTTTTTTTTTTTTTTTTTTTTTTTTTTTTTTACA | | This study |
| | CAGTCTACACATATTCTCTGTTTTTTTTTTTTTTTTTTTTTTTTTTTTTTTTTTTTCCA | | This study |
| | CAGTCTACACATATTCTCTGTTTTTTTTTTTTTTTTTTTTTTTTTTTTTTTTTTTTACC | | This study |

**Supplemental Table S5.** Primers used in this study (Winzeler et al. 1999).

#### Supplemental Table S6

| Plasmid name | Brief Description | Source |
|---|---|---|
| pCR$^{TM}$2.1 | TA cloning vector | Invitrogen K202020 |
| pCS39 | pCR2.1 + 5XUAS landing pad | This study, Shubin, et al 2020 |

**Supplemental Table S6.** Plasmids used in this study.
